# Long non-coding RNA gene regulation and trait associations across human tissues

**DOI:** 10.1101/793091

**Authors:** O. M. de Goede, N. M. Ferraro, D. C. Nachun, A. S. Rao, F. Aguet, A. N. Barbeira, S. E. Castel, S. Kim-Hellmuth, Y. Park, A. J. Scott, B. J. Strober, GTEx Consortium, C. D. Brown, X. Wen, I. M. Hall, A. Battle, T. Lappalainen, H. K. Im, K. G. Ardlie, T. Quertermous, K. Kirkegaard, S. B. Montgomery

## Abstract

Long non-coding RNA (lncRNA) genes are known to have diverse impacts on gene regulation. However, it is still a major challenge to distinguish functional lncRNAs from those that are byproducts of surrounding transcriptional activity. To systematically identify hallmarks of biological function, we used the GTEx v8 data to profile the expression, regulation, network relationships and trait associations of lncRNA genes across 49 tissues encompassing 87 distinct traits. In addition to revealing widespread differences in regulatory patterns between lncRNA and protein-coding genes, we identified novel disease-associated lncRNAs, such as *C6orf3* for psoriasis and *LINC01475*/*RP11-129J12.1* for ulcerative colitis. This work provides a comprehensive resource to interrogate lncRNA genes of interest and annotate cell type and human trait relevance.

**One Sentence Summary:** lncRNA genes have distinctive regulatory patterns and unique trait associations compared to protein-coding genes.

## Main Text

Long non-coding RNA (lncRNA) genes are a prevalent and heterogeneous group of RNA molecules that lack protein-coding potential. They vary in their epigenetic marks, splicing and transcript structure (*1-4*). Previous studies have demonstrated that lncRNA genes have lower expression, increased tissue-specificity, and greater variability in expression across individuals than protein-coding genes (*1, 5-8*). Despite these differences, many lncRNA genes have been demonstrated to have important roles in gene regulation from epigenetic reprogramming to post-transcriptional regulation (*4, 9*). Although the number of annotated lncRNA genes is increasing as a result of more sensitive transcriptomic profiling in a wide range of contexts (*1, 5, 10, 11*), it is not known how many of these lncRNAs have important functional consequences.

In this study, we used the Genotype-Tissue Expression (GTEx) project v8 data to profile genetic regulation of lncRNA genes across 49 human tissues. We combine multiple approaches, including expression quantitative trait loci (eQTL) analysis, gene expression outlier analysis, co-expression networks, and colocalization, to identify putative functional lncRNAs, their cellular contexts, and their relevance to human traits.

### Literature and database curation produced relevant and comparable sets of lncRNA and protein-coding genes

lncRNA genes are difficult to study because of their low expression patterns and heterogeneity. To mitigate this challenge, we incorporated three subgroups of genes into our comparisons (Fig. 1A): protein-coding genes that were expression-matched to lncRNA genes (Fig. S1); high-confidence non-coding lncRNA genes, which passed an especially stringent set of criteria to be classified as non-coding (*12*); and a set of 713 lncRNAs with strong prior evidence of function (*13, 14*) (Table S1) (see Methods). Poly(A) selection was performed prior to RNA-sequencing, which can affect the types of lncRNA genes quantified: RNA-sequencing libraries prepared by ribosomal RNA depletion and by poly(A) selection quantify similar numbers of lncRNA genes, but lncRNA genes unique to poly(A) selection tend to be antisense transcripts, whereas lncRNA genes unique to ribosomal RNA depletion tend to be intergenic or intronic lncRNA genes (*15*).

**Fig. 1.**
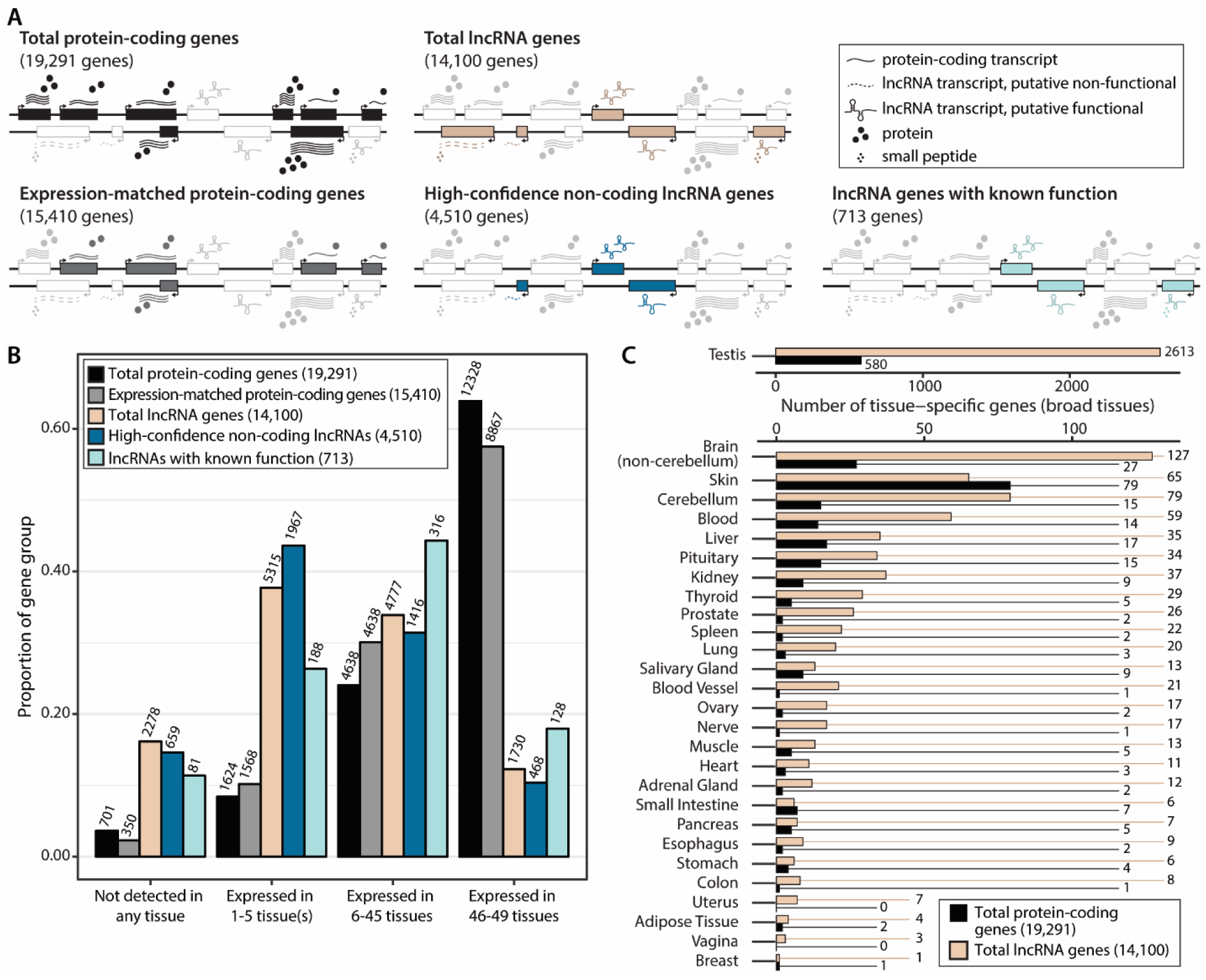
Tissue-specificity of lncRNA and protein-coding gene expression in GTEx. (A) The protein-coding and lncRNA gene groups compared in this paper. The “Expression-matched protein-coding genes” group is a subset of the “Total protein-coding genes”, and the “High-confidence non-coding lncRNA genes” and “lncRNA genes with known function” groups are subsets of the “Total lncRNA genes”. (B) Proportion of each gene group expressed in a certain number of tissues. Bar labels show the number of genes. (C) Numbers of lncRNA and protein-coding genes expressed in only one of the 28 broad tissues. For (B) and (C), the expression threshold is TPM ≥0.5 in >20% of samples.

Before comparing patterns of expression and regulation in these gene groups, we first compared the physical properties of these genes. As has been previously described (*6, 16*), protein-coding genes tended to be longer than lncRNA genes (with median transcript lengths of 3,513 and 657 bases, respectively) and have higher numbers of exons (medians of 10 and 2, respectively) (Fig. S2A-C). Lower proportions of lncRNA genes were expressed in each tissue compared to protein-coding genes, and lncRNA isoforms generally had higher transcript support level scores (Fig. S2D-E). Transcript support level scores reflect the quality of primary data supporting the transcript structure, with lower scores indicating a more well-supported transcript model. The higher scores of lncRNA genes are likely related to the relative scarcity of lncRNA transcripts. Many of these physical differences likely informed the more complex regulatory differences we saw throughout these analyses.

### lncRNA genes have greater tissue-specificity in gene expression and eQTLs

The well-established tissue-specificity of lncRNA gene expression (*1, 5, 6, 8, 11*), was apparent across the 49 analyzed tissues. At an expression threshold of TPM ≥0.5 in at least 20% of samples, most of the genes in both the total lncRNA and the high-confidence non-coding lncRNA gene groups are either not expressed, or only expressed in 1-5 tissues (Fig. 1B). In contrast, the majority of both protein-coding gene groups are expressed in all or nearly all tissues. Expression of lncRNA genes with known function showed intermediate tissue-specificity, with the proportion of genes expressed in only 1-5 tissues significantly lower than total lncRNA genes (χ^2^ = 36.8, p = 1.3 × 10^-9^, test of equal proportions) but significantly greater than total coding genes (χ^2^ = 266.7, p <2.2 × 10^-16^, test of equal proportions). This may reflect the biases in functional lncRNA identification: namely, genes that are operating in many different contexts are more likely to have their functions discovered.

To explore which tissues had the highest numbers of uniquely expressed genes, we focused on the 28 “broad tissue” categories; these are more general tissue types defined by the GTEx Consortium, to which each of the 49 tissues are assigned. There were 4,114 genes that were broad tissue-specific, that is they were expressed in only one broad tissue type (Table S2). Of these uniquely expressed genes, 3,301 were lncRNA genes and only 813 were protein-coding genes (Fig. 1C). Both groups were dominated by genes with testis-specific expression; this may reflect the “transcriptional scanning” suggested to occur during spermatogenesis to reduce the male germline mutation rate (*17*). The next highest numbers of broad tissue-specific genes came from the brain, skin, and blood.

In GTEx v8, 94.7% of protein-coding genes and 67.3% of lncRNA genes were identified as eGenes (i.e. having at least one eQTL at FDR ≤0.05) in at least one tissue (*18*). Per tissue, approximately 85% of expressed protein-coding genes and 50% of expressed lncRNA genes are eGenes (at an expression threshold of TPM ≥0.5 in >20% of samples) (Fig. 2A). The lower frequency of lncRNA eGenes may be partly due to their low gene expression, which limited eQTL detection. However, the fact that approximately 75% of expression-matched protein-coding genes expressed in each tissue were eGenes suggests that there are other factors involved. One such factor could be simpler regulatory mechanisms, which have been reported in certain types of lncRNAs (*19*). With fewer modifiers of their expression, eQTLs for lncRNA genes may occur less frequently, but have stronger effects when present. This is supported by our observation that lead eVariants (the genetic variants with the most significant associations for each gene) for lncRNA genes had larger effect sizes than expression-matched protein-coding genes, with a median log2(allelic fold-change) of 0.805 for lncRNA lead eVariants and 0.579 for expression-matched protein-coding lead eVariants (Wilcoxon rank-sum test W =4.88×10^9^, p <2.2 ×10^-16^; Fig. 2B) (*20*). Higher effect sizes were also reported in a previous study of intergenic long non-coding RNA (lincRNA) eQTLs compared to protein-coding genes, which the authors attributed to less constraint on lincRNA gene expression (*21*). Intriguingly, however, the proportion of eGenes with >1 independent eQTL is comparable between lncRNA and protein-coding eGenes (Fig. 2C).

**Fig. 2.**
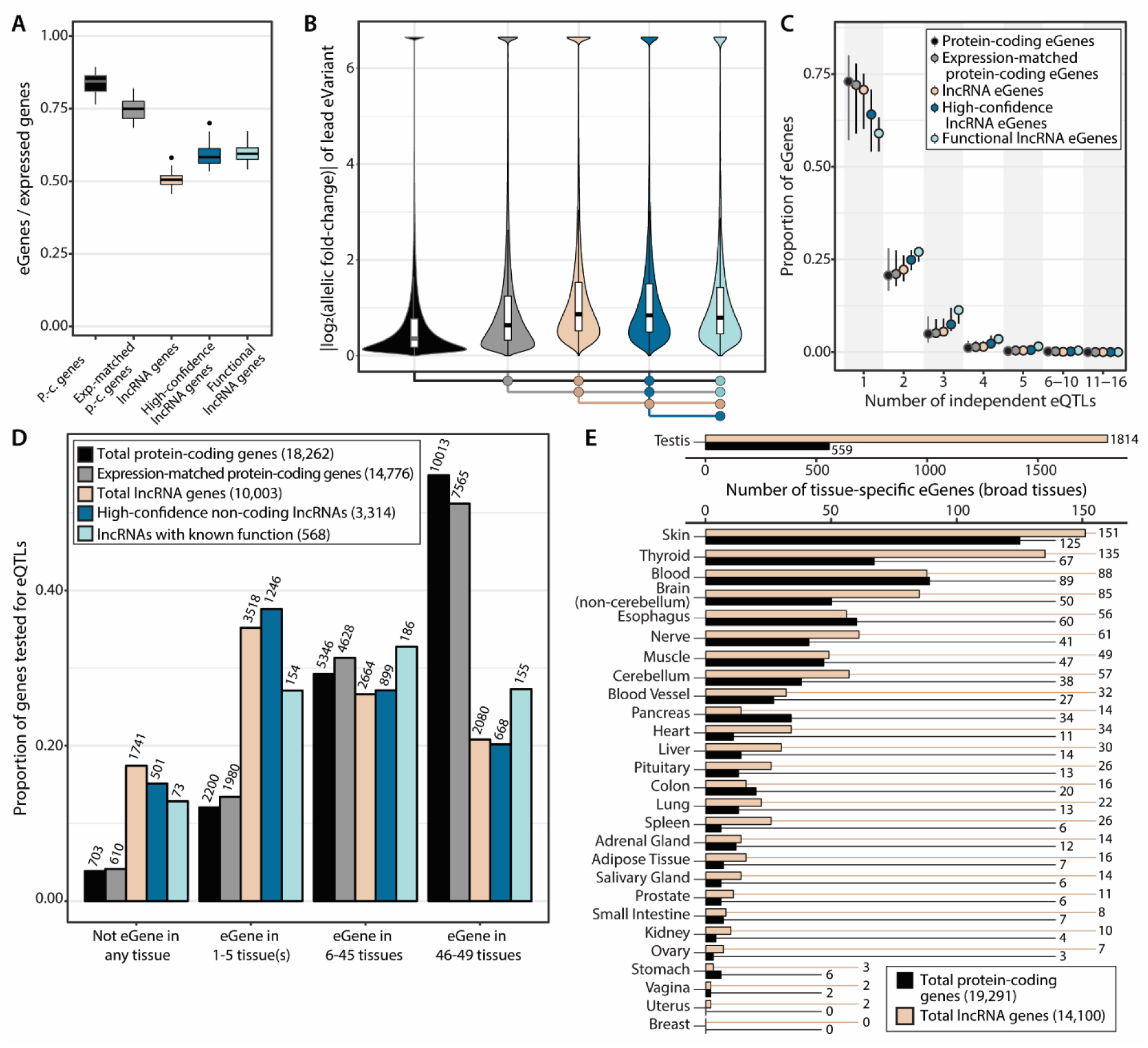
Frequency, effect size, and tissue-specificity of eQTLs in lncRNA genes and protein-coding genes. (A) Proportion of expressed genes that are eGenes (MashR LFSR ≤0.05). Box plots reflect the proportions across the 49 GTEx tissues. (B) Absolute effect size of the most significant eQTL for each gene in each tissue. Effect size is measured as log2(allelic fold-change). at the bottom of the plot indicate significant differences in effect size (Wilcoxon rank-sum test, p-value ≤0.05), with the fill color matching the group with larger eQTL effect size. (C) Distribution of the number of independent eQTLs for eGenes in each gene group. Circles represent the median proportion of eGenes across the 49 GTEx tissues with that number of independent eQTLs, and whiskers extend the interquartile range. (D) Proportion of each gene group that is an eGene in a certain number of tissues. Bar labels show the number of genes. Total numbers in the legend reflect the number of genes that were tested for an eQTL in at least one tissue, and thus differ from the numbers in Figure 1D. (E) Numbers of lncRNA and protein-coding eGenes specific to one of the 28 broad tissues. For (A), (B), (D) and (E), eQTLs were identified using the MashR method with a significance threshold of LFSR ≤0.05. For (C), independent eQTLs were identified using the forward stepwise regression-backwards selection method (see Methods). p.-c. genes = protein-coding genes.

The tissue-specificity of eGenes across all gene groups (Fig. 2D) follows the patterns of tissue-specificity in gene expression (Fig. 1D), with lncRNA eGenes generally observed in fewer tissues than protein-coding eGenes. There were more broad tissue-specific eGenes than there were genes with broad tissue-specific expression: 2,783 lncRNA eGenes, and 1,267 protein-coding eGenes (Fig. 2E; Table S3). Given the similar patterns of tissue-specificity in both gene expression and the presence of eGenes, a natural assumption would be that many of the tissue-specific eGenes are simply genes with tissue-specific expression that have eQTLs. Surprisingly, this was rarely the case: except for the testis-specific eGenes and prostate-specific lncRNA eGenes, fewer than 25% of tissue-specific eGenes also showed tissue-specific gene expression (median overlap 4.5%) (Fig. S3).

Previous studies have found that different subtypes of lncRNA genes have different promoter structure and expression patterns (*1, 19*). Of the GENCODE lncRNA biotypes, we noted that 62% of lncRNAs with tissue-specific expression were lincRNAs, which was higher than their proportion in total lncRNAs (53%; χ^2^ = 142.5, p <2.2 ×10^-16^, test of equal proportions) (Fig. S4A). In contrast, antisense lncRNAs were depleted for tissue-specific expression: 31% of lncRNA genes with tissue-specific expression were antisense, compared to 37% of the total set (χ^2^ = 66.2, p = 4.1 ×10^-16^, test of equal proportions). This is consistent with previous reports of lincRNAs having notably high tissue-specificity in comparison to lncRNA genes that diverge from another gene (via bidirectional transcription) (*19*), since many of the GENCODE-annotated antisense lncRNA genes are identified to actually be promoter-divergent in the FANTOM CAGE-associated transcriptome (FANTOM-CAT) (*1*) (Fig. S4B). It was also interesting to note that 15% of GENCODE intergenic lncRNAs were identified as promoter-divergent in FANTOM-CAT. This highlights the importance of maintaining updated gene annotations, especially when examining the frequently updated lncRNA genetic landscape (*22*).

### Multi-tissue outliers for intergenic lncRNA gene expression are frequently overexpressed

Given that lncRNA regulation is often tissue-specific, we were interested in whether individuals can defy these patterns and display outlier lncRNA gene expression in non-canonical tissues. To test this, we confined the outlier gene expression analysis performed in the GTEx rare variants paper (*23*) to only examine lincRNA genes, the lncRNA gene subtype that most commonly displayed tissue-specific expression (Fig. S4A). To identify multi-tissue outliers, we also limited our analysis to individuals with expression data available for the given gene in at least 5 tissues. Overall, 2,535 individual-lincRNA combinations were found to be expressed at more than 2 standard deviations above or below the mean in a majority of tested tissues; these were termed multi-tissue outliers (|median Z-score| > 2) (Table S4). These outlier events (with an event being an individual-lincRNA combination) involved 1,009 unique lincRNAs out of the 4,236 tested. The majority of lincRNA outliers (86%) have a positive median Z-score, which means that the outlier individuals typically over-expressed that lincRNA gene (Fig. 3A). For protein-coding genes, only 61% of outliers at the same threshold are over-expressed (*23*). This is mostly attributable to the lower expression of lincRNA genes compared to protein-coding genes; lowly expressed genes are more likely to fluctuate upward, and their under-expression is difficult to detect.

**Fig. 3.**
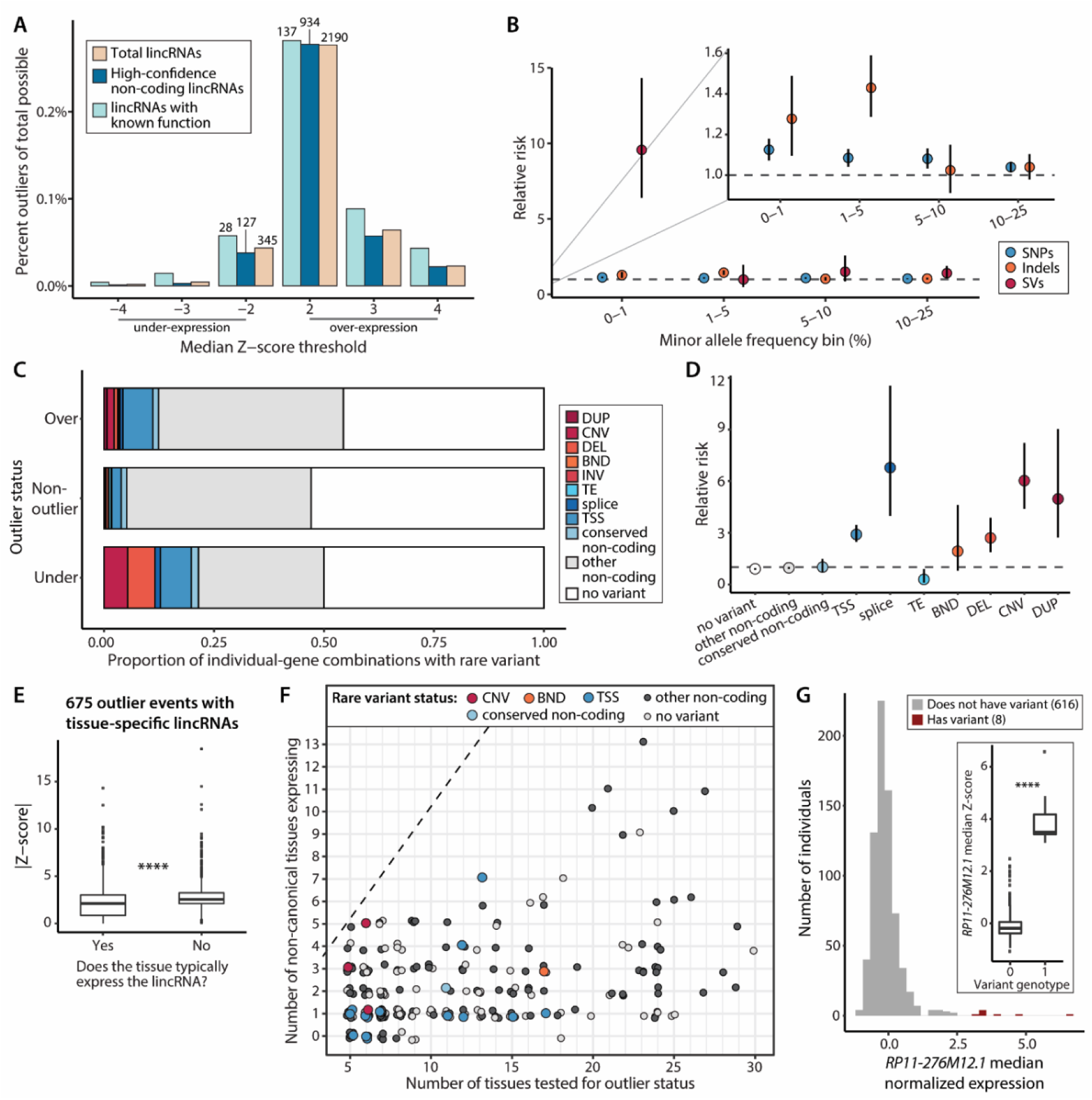
Outliers in lincRNA expression. (A) Percent of multi-tissue lincRNA gene outliers (gene-individual combinations) out of all gene-individual combinations tested. Labels indicate the number of outliers. (B) Enrichment of variants within 10kb of the outlier gene in outlier individuals. (C) Presence of rare variants (MAF <1%) within 10kb of the outlier gene based on outlier status. (D) Enrichment of rare variants (MAF <1%) within 10 kb of the outlier gene in outlier individuals. (E) Tissue-specific Z-scores for outlier events involving lincRNAs with tissue-specific gene expression (as identified in Fig. 1), separated by whether or not the tissue typically expresses that lincRNA. (F) For outlier events involving tissue-specific lincRNA genes, the number of non-canonical tissues expressing the gene in the outlier individual versus the number of their tissues that were tested for outlier status. Non-canonical expression was TPM ≥0.5 in the outlier individual’s sample, for a tissue had TPM <0.1 in >80% of samples. Calling non-canonical expression was not limited to the tissues that were tested for outliers (so an individual could have more tissues with non-canonical expression than tissues tested). (G) Normalized expression of *RP11-276M12.1*, a gene that was a multi-tissue outlier for 13 individuals. Each value on the histogram is one individual’s median expression of the gene across all tissues. Of these individuals, 8 had the same rare variant in the first exon (filled in red). Inset: Individuals’ median Z-scores for this gene separated by presence or absence of the rare variant. DUP = duplication, CNV = copy number variation, DEL = deletion, BND = breakend, TE = transposable element insertion, TSS = transcription splice site.

For each outlier, we identified variants within 10kb of the outlier gene for individuals with self-reported European ancestry, as allele frequencies may not be consistent across populations (*23*). Outlier individuals were more enriched for nearby variants with lower allele frequencies (MAF < 1%; see Methods), with relative risks (RRs) of 1.12 for SNPs, 1.28 for indels, and 9.56 for SVs (Fig. 3B). In both over- and under-expression outliers, higher proportions of nearby rare variants were observed compared to non-outliers (54.4% of over-expression outliers, 50% of under-expression outliers, and 47% of non-outliers), though the proportion of overall rare variants nearby under-expression outliers vs non-outliers was not significantly enriched (RR over = 1.16, p = 2.82 × 10^-9^, RR under = 1.06, p = 0.37, Wald test). Rare structural variants were more often associated with under-expression of the lincRNA (RR = 10.53, p = 5.59 × 10^-20^, Wald test), and rare variants in transcription start sites (TSS) were enriched in both directions (Fig. 3C). Of the rare structural variants driving the enrichment nearby outliers (Fig. 3B), we found that deletions, CNVs, and duplications were specifically enriched in outlier individuals near their outlier genes (Fig. 3D). However, rare splice variants were also strongly enriched nearby outlier genes (RR = 6.78, p = 5.20 × 10^-8^) - even more so than rare TSS variants (RR = 2.92, p = 4.35 × 10^-26^). This indicates that transcript structure is a key influence on overall expression; splicing variation may mediate this by affecting transcript maintenance and decay. This is also supported by the similarly strong enrichment for rare splice variants observed near protein-coding gene outliers (*23*), as well as the strong enrichment of splice-related annotations for *cis*-eQTLs, including those that were distinct from splice QTLs (*18*).

We next investigated how many multi-tissue outliers disrupted the tissue-specific patterns of lincRNA gene expression. Of the 2,535 outliers, 675 involve lincRNAs that were only expressed in 1-5 of the 49 GTEx tissues (301 unique genes). Per-tissue Z-scores were lower in the tissues that typically expressed these outlier genes compared to those that did not (median in expressing tissues = 2.1, median in non-expressing tissues = 2.5, Wilcoxon rank-sum test p <2.2e-16) (Fig. 3E). This suggests that outlier events involving tissue-specific lincRNAs could have particularly dramatic effects by involving aberrant expression in tissues that do not usually express the lincRNA. This occurred in 296 outlier events in which lincRNAs with tissue-specific expression had TPM ≥0.5 in at least one non-canonical tissue (in which the lincRNA gene’s TPM <0.1 in >80% of samples) (Fig. 3F).

One notable outlier involves the gene *RP11-276M12.1* (ENSG00000259445). This lincRNA gene was typically expressed in three tissues: thyroid, testis, and vagina. There were 13 over-expression outlier individuals for this gene, of whom 12 were assessed for the presence of rare variants. Of these 12 individuals, 11 had rare variants nearby the gene, and 8 of these individuals had the same rare variant located in the first exon, within 100 bases of the TSS (Fig. 3G). *RP11-276M12.1* was expressed at TPM ≥0.5 in 0 to 4 non-canonical tissues, depending on the outlier individual (median =1). This was spread across eight different tissues, with tibial artery showing non-canonical expression of the gene in eight outlier individuals. The individual who expressed *RP11-276M12.1* at TPM ≥0.5 in four different non-canonical tissues did so in the two cerebellum tissues, spinal cord (cervical-C1), and cultured fibroblast cells. This gene is interesting not just because of the high number of outlier individuals, but also that many of them share the same rare variant. This variant, a G>C substitution at chromosome 15 position 81,995,598 (hg38 assembly), has a frequency of 0.0063 in GTEx and 0.0066 in gnomAD non-Finnish Europeans (0.0045 overall). It is associated with a significant difference in expression (Wilcoxon rank-sum test p =1.2e-06; Fig. 3G), but was missed by traditional eQTL testing that filters out variants with MAF <0.01.

### eVariants can be shared between lncRNA and protein-coding genes

Since many lncRNA genes have *cis-*regulatory effects on other nearby genes (*24*), we assessed the prevalence of shared genetic effects. Across all independent lead eVariants, 7.5% were associated with >1 gene in the same tissue (Fig. 4A). The other 92.5% were only associated with 1 gene, most of which were protein-coding genes (65.2%, compared to 17.3% for lncRNAs and 10.0% for all other gene types).

**Fig. 4.**
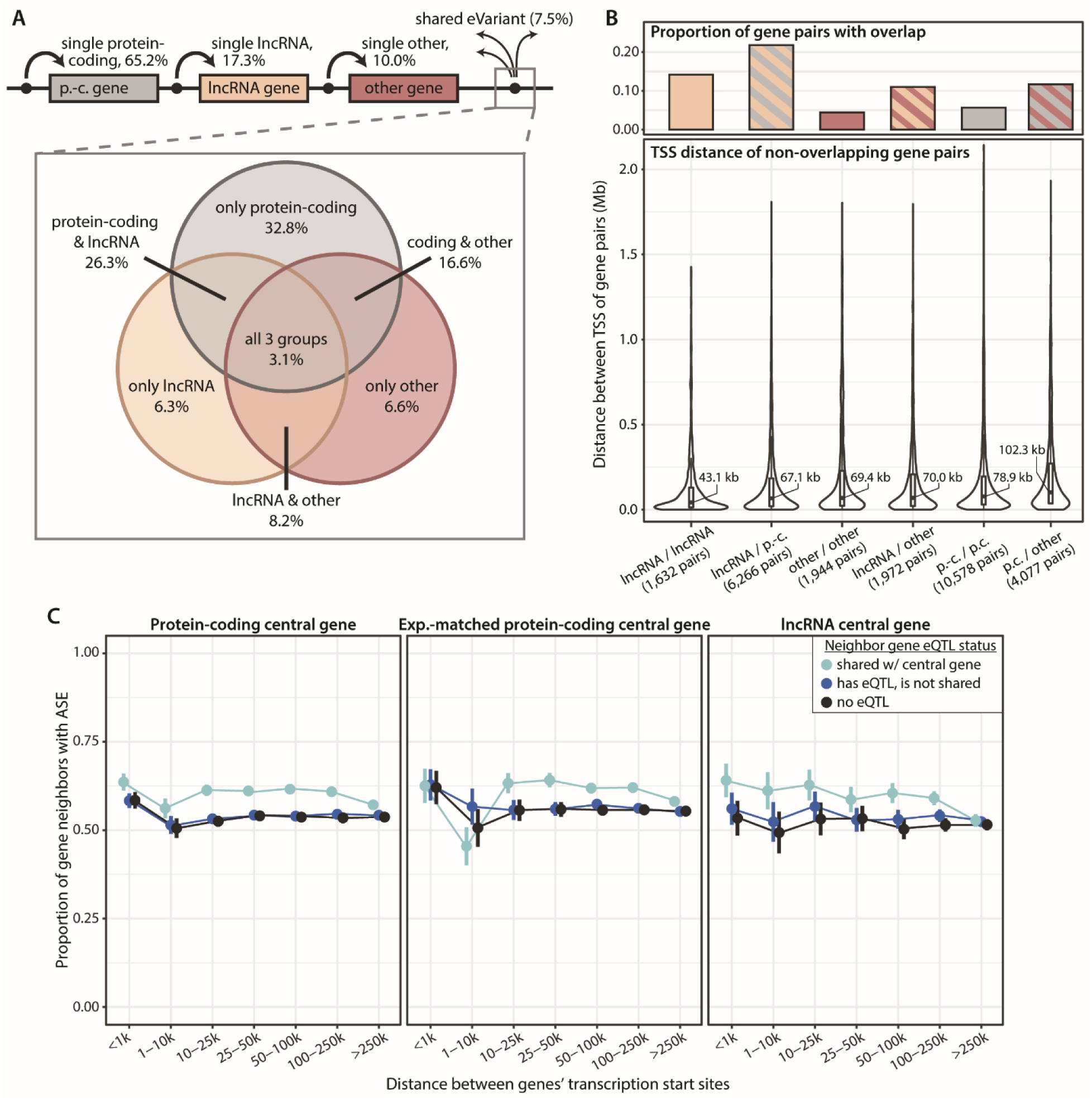
Connecting genes through shared eVariants. (A) Summary of eVariant sharing using the independent eQTLs (see Methods). Genes shared an eVariant if their expression was associated with either the same eVariant, or with eVariants that were within 500kb of each other with an R2 ≥0.85, in the same tissue. *(top)* percentage of eQTLs only associated with one gene (“single”) versus how many were associated with >1 gene (“shared”). *(bottom)* The shared eVariants categorized based on which types of genes they were associated with. (B) *(top)* Proportion of eVariant-sharing gene pairs that have overlapping genomic location. *(bottom)* Distance between TSS’s of non-overlapping gene pairs that share an eVariant. (C) Extension of allele-specific expression (ASE) from a central gene with strong ASE (adjusted binomial p-value ≤0.05, allelic ratio deviation from 0.5 of 0.35-0.48). The line colors indicate whether the neighboring gene has an eQTL, and whether its associated variant is shared with the central gene. Neighboring genes were limited to non-overlapping protein-coding genes only, and were checked for significant ASE (adjusted binomial p-value ≤0.05). Bars indicate the 95% CI across different central genes, with ASE status being collapsed by individuals and tissues. p.-c. gene = protein-coding gene.

When the shared eVariants were broken down by which types of genes shared them, a notable proportion (26.3%) were shared between protein-coding and lncRNA genes, which was second only to the proportion shared between multiple protein-coding genes (32.8%). This high occurrence of shared eVariants with lncRNA genes may be partly driven by both antisense-sense gene pairs (13.4% of protein-coding-lncRNA gene pairs that share an eVariant are antisense-sense gene pairs), *cis*-regulatory relationships or shorter distances between lncRNA and protein-coding gene pairs (Fig. 4B).

Of the 26,469 gene pairs sharing eVariants in at least one tissue, 3,568 (13.5%) had cross-mappability scores greater than zero (*24*). Higher cross-mappability scores between gene pairs indicate a greater amount of sequence similarity, and thus greater potential for incorrect read alignment to have affected quantification. Our observed percentage of cross-mapping pairs is higher than the 2.45-4.92% of evaluated gene pairs in the GTEx v7 dataset that were reported as cross-mappable (*25*), suggesting that some of the eVariant sharing may actually be due to cross-mapping. Although the eVariant-sharing gene pair types involving lncRNA genes actually have the lowest proportions of cross-mappable gene pairs (Fig. S5A), this is still an important technical factor to consider; as such, we reported the symmetric cross-mappability score results for all eVariant-sharing gene pairs (Table S5).

Several lncRNA genes shared eVariants across multiple tissues. One such gene was *KANSL1-AS1* (ENSG00000214401), which was connected to 5 different protein-coding genes across many tissues, with the genes sometimes even sharing more than one eVariant (Fig. S5B). These recurring connections may highlight key sets of co-regulated genes.

### Most *cis*-regulation by lncRNA genes operates on a local scale

One of the best-known lncRNA genes, *XIST*, operates in *cis* on a massive scale, inhibiting almost the entire X-chromosome from which its expressed (*26, 27*). We were curious about other lncRNA genes’ range of local regulation, which we assessed via allele-specific expression (ASE). We defined genes with strong ASE (multiple test-adjusted binomial p-value ≤0.05 and allele ratio either 0.02-0.15 or 0.85-0.98 for any variant in the gene) as “central genes”, and then tracked how often significant ASE (multiple test-adjusted binomial p-value ≤0.05, no allele ratio threshold) was maintained in the same tissue in non-overlapping, protein-coding gene neighbors.

Extending out from both protein-coding and lncRNA central genes, over half of all non-overlapping protein-coding gene neighbors also showed significant ASE, with the degree of sharing decreasing as the distance between the genes increased. Nearby downstream neighbors of lncRNA genes had the highest proportions of ASE, and the drop-off in ASE maintenance with distance was greatest from lncRNA central genes (Fig S6A), potentially reflecting antisense-sense gene pair relationships or other mechanisms for *cis*-effects, such as localized epigenetic changes.

Pairs of genes that shared eVariants were more likely to share ASE, compared to central genes and gene neighbors with unshared eQTLs (χ^2^ = 157.7 for lncRNA central genes and 403.4 for protein-coding central genes, both p <2.2 × 10^-16^, test of equal proportions) (Fig. 4C). For lncRNA central genes, this relationship appeared to be dependent on distance. Altogether, these patterns of ASE suggest that most lncRNA genes operate differently than *XIST*, and tend to affect the genes immediately around themselves, if at all, rather than have more far-reaching effects. One interesting example of local ASE was with the lncRNA gene *RP4-568C11.4* (ENSG00000274173), in which 52/59 individuals also displayed ASE in two nearby protein-coding genes (Fig. S6B). Remarkably, for all of these individuals the significant ASE occurred in the same tissue (whole blood).

Despite these overall patterns, there were some examples of lncRNA genes surrounded by large regions of ASE. One such example was the central lncRNA gene *MIR210HG* (ENSG00000247095), for which ASE was seen in protein-coding genes over a range of 358kb downstream to 444kb upstream of the gene in 48 individuals (Fig. S6C).

### Co-expression networks identify highly connected lncRNA genes with specialized cell type associations

For each tissue, we built co-expression networks using weighted correlation gene network analysis (WGCNA) (*28*). Since the goal was to identify lncRNA genes with cell type associations, each module was annotated based on enrichment for gene sets associated with Gene Ontology Cellular Compartments and cell types from blood, the central nervous system, and the Mouse Cell Atlas (*29*) (see Methods). The number of modules created per tissue was not associated with the sample size of the tissue (Fig. S7A). A median of 48% of modules in each tissue were annotated, ranging from 18% of modules in the ovary to 85% of modules in the stomach (Fig. S7B).

With the exception of testis tissue, approximately 50% of lncRNAs did not meet the expression requirements to be included in the co-expression networks (median of excluded lncRNA genes across tissues = 47%) (Fig. 5A). The proportion of excluded genes was lower for lncRNA genes with known function (median = 35%). Greater proportions of protein-coding genes (both the total group and the expression-matched group) were assigned to modules compared to lncRNA genes (Fig. 5A). Generally, larger modules included higher proportions of lncRNA genes (Fig. 5B); however, there were also some smaller modules mostly made up of lncRNA genes, which may be worthy of further exploration as potential hubs of lncRNA regulatory activity in those tissues.

**Fig. 5.**
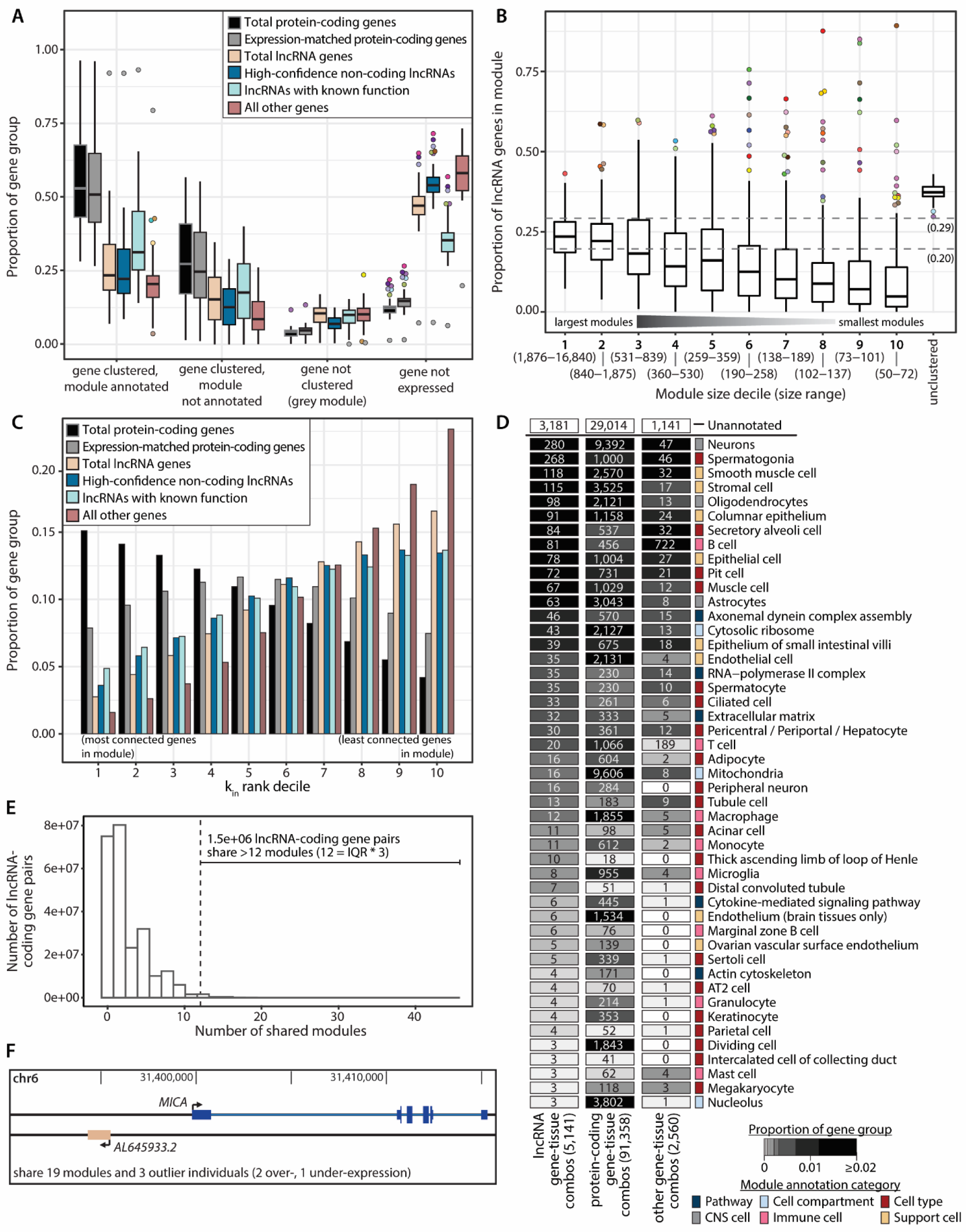
Connecting genes through weighted gene co-expression network analysis (WGCNA). (A) Summary of gene assignment to modules by gene group and tissue. The underlying box plot shows the proportion of a gene group falling into that module status across tissues. Outlier point color indicates the tissue. (B) Proportion of lncRNA genes in modules across all tissues, binned by module size. Horizontal dashed lines indicate the highest and lowest proportions of lncRNA genes that met the expression threshold for WGCNA across all tissues (0.29 = testis, 0.20 = whole blood). (C) Proportion of gene groups by intra-modular connectivity (kin) ranking. The most-connected genes within their module are in the first kin rank decile, and the least-connected genes within their module are in the tenth kin rank decile. (D) Module annotations of genes with high intra-modular connectivity (the gene is in the top kin rank decile of its module, and has scaled kin ≥0.5). Box fill reflects the proportion of genes assigned to a module with that annotation. Since genes are assigned to modules in multiple tissues, the labels reflect gene-tissue combinations, not individual genes. (E) Distribution of the number of modules shared between unique lncRNA/protein-coding gene pairs. (F) Location of the protein-coding gene *MICA* and the lncRNA gene *AL645933.2*, which share modules in 19 tissues and also are both outlier genes in 3 individuals.

We evaluated how strongly connected genes were based on their intramodular connectivity (k_in_) values, which were converted to a rank and grouped into deciles within each module. Higher proportions of protein-coding genes were among the highest-ranked (most connected) genes, although this was expression-dependent (Fig. 5C). The majority of lncRNA genes in each module were not highly connected, indicating that most lncRNAs only have potential regulatory relationships with one or few nearby genes. However, although they were in the minority, there were also some highly connected lncRNA genes that may represent lncRNAs with potentially far-reaching regulatory effects (Table S6). We also incorporated an “all other gene types” group, which included all non-coding genes that are not lncRNAs as well as pseudogenes. Since many of the quantifications in this group are of poor quality, it served as a measure of noise in the networks. As expected, lncRNAs were generally better connected than this “all other gene types” category (Fig. 5C; p <2.2 × 10^-16^, Wilcoxon rank-sum test of decile rankings between the “total lncRNA genes” group and “all other genes” group). With module membership values, which reflect how well a gene’s expression correlates with its assigned module, the same trend was observed of protein-coding genes having the highest scores and lncRNA genes having lower module membership, but not as low as the catch-all “all other gene types” category (Fig. S7C).

We next examined the most common annotations of highly connected genes. To be considered “highly connected”, a gene had to be within the top decile of its module based on kin values and have a scaled kin value ≥0.5. Of the highly connected lncRNAs (5,141 gene-tissue combinations), only 1,960 were assigned to annotated modules (Fig. 5D). The high proportion of well-connected lncRNA genes in unannotated modules suggests that many of the pathways involving lncRNAs have yet to be identified.

Compared to the modules that protein-coding genes were assigned to, a higher proportion of lncRNA annotation terms related to specialized cell types such as secretory alveoli cells of lungs, B cells of the immune system, pit cells of the liver, spermatocytes, and kidney regions like tubule cell, loop of Henle, and distal convoluted tubule (Fig. 5D, Fig. S7D). In contrast, protein-coding genes were more frequently assigned to modules annotated with common cell types and ubiquitous cell compartments terms such as endothelium, resident macrophages, mitochondria, and nucleolus. Although lncRNA genes rarely formed the hubs of a given module, the annotation of these networks provides a resource to identify cell type or compartment-related lncRNA genes.

There were >1.5 × 10^6^ unique lncRNA/protein-coding gene pairs that shared modules in >12 tissues, which was 3*IQR of module sharing across all lncRNA-coding gene pairs (Fig. 5E; Table S7). 614 of these gene pairs also shared at least 1 multi-tissue outlier individual for both genes (Fig. S7E). Of these 614 co-expressed genes with shared outliers, 71 of them involved a rare variant within 10kb of the lncRNA gene (Fig. S7F). These 614 gene pairs are compelling candidates to explore further for functional links.

There were some notable candidate lncRNA/protein-coding gene pairs due to a high degree of module sharing (N >12) and multiple outlier events. The first pair, *MICA* and *AL645933.2* (ENSG00000272221), were both over-expression outliers in two individuals, and both under-expression outliers in one individual. One of these individuals had different rare SNVs within 10kb of both genes (Fig. 5F). *MICA* encodes major histocompatability complex class I chain-related protein A, a highly polymorphic stress-induced antigen that is associated with inflammatory responses and cancer (*30-32*). The co-expression of *AL645933.2* with *MICA* suggests that it may also have a role in these processes, although 13 of the modules they share across 19 tissues are unannotated. Another gene pair, *ACOT1* and *AC007228.9* (ENSG00000268568), was intriguing because they are present on separate chromosomes. These genes shared two under-expression outlier individuals. In both cases, the individual had a rare SNV within 10kb of *AC007228.9. ACOT1*, which encodes acyl-CoA thioesterase 1, is involved in lipid metabolism. Given that these two genes’ shared modules were frequently annotated as mitochondria modules, as well as being assigned to the adipocyte module in the adipose (visceral omentum) tissue, *AC007228.9* may also be involved in lipid metabolism.

### The majority of significant lncRNA colocalization events have a stronger signal than nearby protein-coding genes

Colocalization analyses connect genetic variation, gene expression, and traits by integrating results from eQTL analyses and genome-wide association studies (GWAS). As part of the main GTEx paper (*18*) and GWAS companion paper (*33*), colocalization analyses were performed for 87 traits in each tissue. There were 4,694 significant events (posterior probability of a shared predicted causal variant between eQTL and GWAS (PP4) ≥0.5) involving lncRNA genes, which encompassed 48 traits and 690 lncRNA genes (Table S8). As a point of reference, there were 20,281 significant protein-coding gene colocalization events, involving 53 traits and 2,785 unique genes.

For most of the lncRNA gene colocalization events (3,757, 80.0%), there was no significant protein-coding gene colocalization within 500kb up or downstream of the lncRNA gene (Fig. 6A). This is not due to an absence of in-range protein-coding genes; there were only 32 events (0.7%) in which there was no annotated protein-coding gene within this range, and those events were excluded from Fig. 6A. Of the lncRNA colocalizations that also had a significant in-range protein-coding gene colocalization, most pairs had similar PP4 values (479, 10.2%), followed by a nearly even split between events with a notably higher protein-coding gene PP4 (239, 5.1%) and ones with a notably higher lncRNA gene PP4 (187, 4.0%). Compared to the total set of lncRNA genes, lincRNA genes alone had an even greater proportion of colocalization events with no in-range significant protein-coding gene, reflecting their independent regulation. In contrast, fewer colocalization events involving antisense lncRNAs had no in-range significant protein-coding genes.

**Fig. 6.**
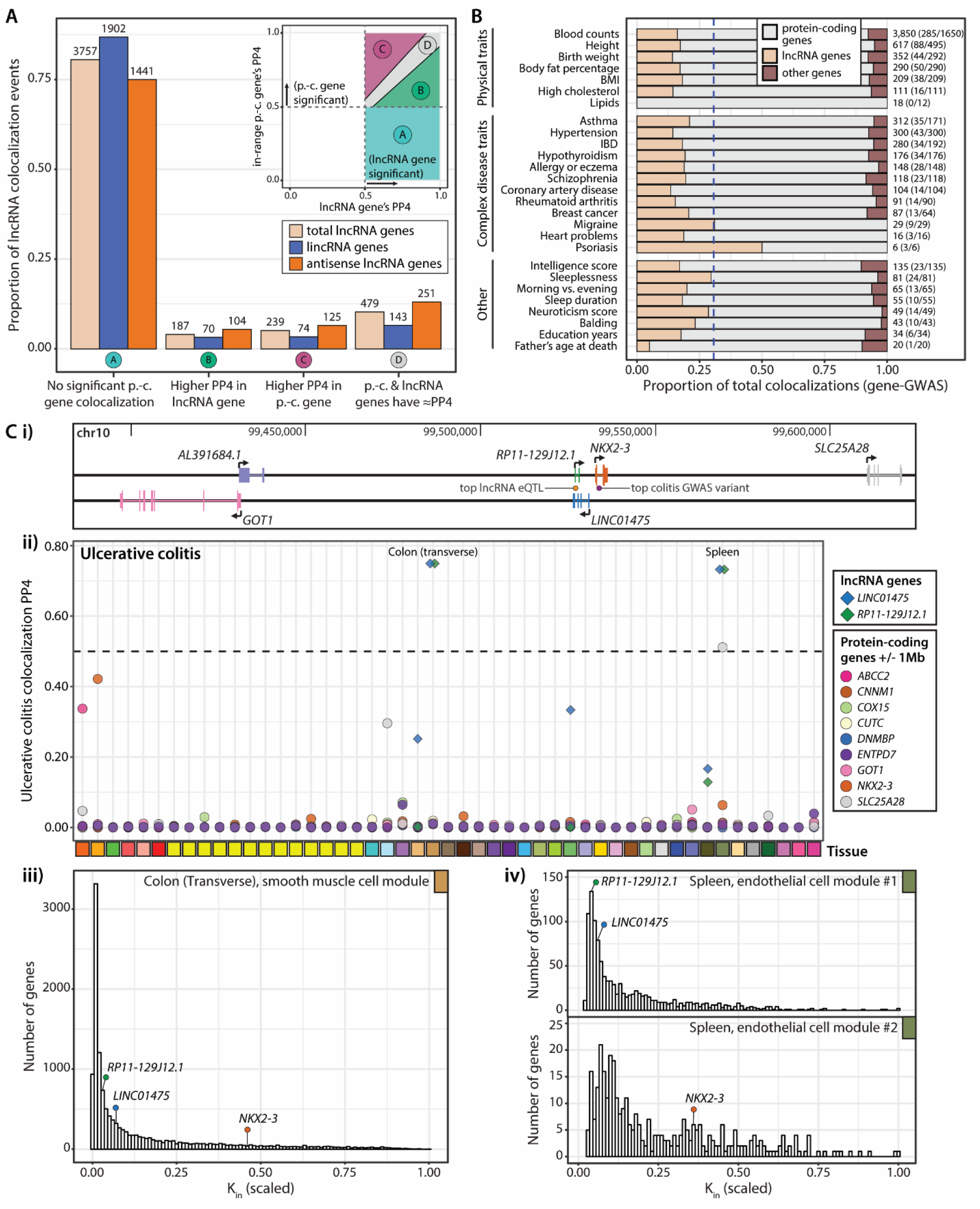
GWAS-eQTL colocalization events involving lncRNA genes. (A) Summary of significant lncRNA colocalization events (event = gene-tissue-GWAS combination) by their relationship to the protein-coding gene with the highest PP4 within 1Mb of the significant lncRNA gene. There were 32 significant lncRNA colocalization events where there was no protein-coding gene in range, which are not included in this plot. (B) Contribution of each gene type to significant colocalization events, collapsed across tissues (i.e. gene-GWAS combinations). GWASes were grouped on the y-axis by more general trait categories, and any traits with fewer than 5 significant gene-GWAS combinations were not included in this plot. The dashed line is the proportion of genes tested for colocalization that were lncRNAs (0.31). The numbers to the right of each bar show the total number of significant colocalization events (gene-trait-tissue combinations), followed by the number of unique significant lncRNA genes / the total number of unique significant genes in brackets. (C) Exemplar significant colocalizations with *LINC01475* and *RP11-129J12.1* and ulcerative colitis. i) Location of the lncRNA genes, as well as nearby protein-coding genes. Locations of the most significant ulcerative colitis GWAS variant and the top eQTL for both lncRNA genes in the transverse colon are indicated. ii) Colocalization posterior probability values across each tissue for *LINC01475* and *RP11-129J12.1*, as well as all protein-coding genes within 1Mb of the lncRNA genes, with ulcerative colitis. The dashed line indicates the threshold for significance, PP4 ≥0.5. iii) Scaled intramodular connectivity (kin) of *LINC01475, RP11-129J12.1*, and *NKX2-3* within their assigned smooth muscle cell module in the transverse colon gene co-expression network. iv) Scaled intramodular connectivity (kin) of *LINC01475, RP11-129J12.1*, and *NKX2-3* within their assigned endothelial cell modules (two separate modules) of the spleen gene co-expression network.

Across all significant colocalization events, lncRNA genes were depleted compared to their proportion in the gene set tested (Fig. 6B). However, the proportion of lncRNA colocalization events varied by trait type, with the highest percentages occurring in migraine (31%), and psoriasis (50%). For psoriasis, colocalization events were dominated by one lncRNA gene. The single-exon lncRNA *C6orf3* (ENSG00000255389), which runs sense intronic to the protein-coding gene *TRAF3IP2*, showed significant colocalization in 35 tissues. No protein-coding genes in the 1Mb neighborhood around *C6orf3* had a significant colocalization with psoriasis in any tissue (Fig. S8A). The tissue with the highest PP4 value (0.996) was sun-exposed skin, which is compelling given that psoriasis is a chronic skin condition. Notably, top significant eQTLs for *C6orf3* are the exact top GWAS variants, which does not always occur with colocalization (Fig. S8B). In both skin tissues (sun-exposed and not sun-exposed), *C6orf3* clustered in the “endothelial cell”-annotated co-expression module (Fig. S7C).

For significant lncRNA colocalizations, the in-range protein-coding gene with the highest PP4 was not always the closest gene (Fig. S8A). This indicates that many of these connections (i.e. the sharing of a significant colocalization) were not just the result of proximity, but may also reflect some regulatory relationship. For the events in which there was no in-range protein-coding gene with a significant colocalization, this connection was essentially arbitrary, since the protein-coding genes’ PP4 was <0.5 in these cases. Thus, these lncRNA/protein-coding gene pairings have limited relevance, which is reflected in the wide range of distances between the two genes (Fig. S8A). It is also worth noting that, for 1,435 of the 3,757 events with no in-range significant protein-coding gene, the significant lncRNA gene actually overlapped a protein-coding gene, but it still did not have a significant colocalization. For example, this was the case with *C6orf3* and *TRAF3IP2*.

Another interesting case involved the lncRNA genes *LINC01475* (ENSG00000257582) and *RP11-129J12.1* (ENSG00000228778) and the protein-coding gene *NKX2-3* in relation to ulcerative colitis. The two lncRNA genes have antisense overlap with each other, and are just upstream from *NKX2-3* (Fig. 6Ci). *NKX2-3* has received attention related to the significant colitis GWAS results in this genomic region (*34*). However, it did not have a significant colocalization in any tissue, whereas the two lncRNA genes had the best colocalizations in the transverse colon (PP4 = 0.749 for both genes) and the spleen (PP4 = 0.732 for both genes) (Fig. 6Cii, Fig. S9B).

Both of the tissues in which colocalization occurred are logical for ulcerative colitis: the colon has an obvious role, and the spleen could be connected via immune system regulation. In the co-expression network for the transverse colon, all three genes were assigned to the same “smooth muscle cell” module, which was unsurprising given their correlated gene expression (Fig. S9C). In the spleen co-expression network, *NKX2-3* was assigned to one “endothelial cell” module, and the two lncRNA genes were assigned to a different “endothelial cell” module (Fig. 6Ciii-iv). *NKX2-3* is a homeobox gene that is key for the development of the spleen and the visceral mesoderm, which develops several essential cell types of the gastrointestinal tract including endothelial cells, immune cells, and - notably - smooth muscle cells. Knockout mouse studies have shown that loss of this gene affects spleen architecture, and lymphocyte maturation and homing (*35-39*). These findings make a compelling case that lncRNA regulation of *NKX2-3* in both the colon and spleen influences ulcerative colitis susceptibility.

## Discussion

lncRNA genes differed from protein-coding genes at nearly all levels of regulation. Compared to protein-coding genes, lncRNAs showed greater tissue-specificity in both expression and presence of eQTLs, with the latter not entirely attributable to differences in expression levels, as well as lower intramodular connectivity in gene co-expression networks. Setting lncRNA genes apart even more was the striking number of significant lncRNA colocalization events where there was no significant protein-coding colocalization within 1 Mb (3,757/4,694, 80%). There were often differences between lncRNA subtypes as well: intergenic lncRNA genes more frequently showed tissue-specific expression and significant colocalization events with no nearby significant protein-coding gene colocalizations, whereas antisense lncRNA genes more frequently shared eVariants with other genes. These subtype-specific trends show the importance of maintaining updated lncRNA gene annotation, and interpreting them cautiously. For instance, many lncRNA subtype assignments did not coincide between FANTOM-CAT and GENCODE (*1*). In the current version of GENCODE (v31), they have dispensed with subtype categorization entirely, referring to them all as gene type ‘lncRNA’. It is clear that different subtypes of lncRNA genes have different regulatory patterns and perhaps different roles, and subtype should be considered in any analysis of these genes.

Not only do these analyses highlight the differences between lncRNA and protein-coding genes, but they can also be used to interrogate lncRNA genes of interest and systematically identify lncRNAs associated with certain cell types or traits. There are other resources that provide lncRNA gene networks, conservation data, or expression-based assessments (*11, 40-43*); our analyses examine several of these characteristics, and also provide trait association information. Searching for convergence of evidence across multiple lncRNA resources here enabled identification of the most compelling candidate genes for further study.

Although exploring non-coding genetic variation has become increasingly important, efforts to date have mostly focused on regulatory effects on protein-coding genes. This work provides an important pathway to enhance these efforts towards evaluating non-coding genes and their roles in complex traits and diseases.

## Acknowledgments

We thank the Montgomery and Kirkegaard labs for their feedback on this work.

## Author contributions

O.M.de G. co-led manuscript, conducted analyses, visualized data, and co-wrote manuscript; N.M.F. conducted outlier analysis and contributed to writing; D.C.N. conducted network analysis; A.S.R. contributed to colocalization analysis; F.A. generated QTL and ASE data, and provided feedback on manuscript; A.N.B. contributed to GWAS harmonization and colocalization analysis; S.E.C. generated ASE and tissue sharing (MashR) data and provided feedback on manuscript; S.K.-H. generated tissue sharing (MashR) data and provided feedback on manuscript; Y.P. contributed to colocalization analysis and provided feedback on manuscript; A.J.S. generated structural variant data and provided feedback on manuscript; B.J.S. contributed to outlier analysis; A.B. contributed to outlier analysis and provided feedback on manuscript; C.D.B. led trainees and contributed to GWAS analysis; X.W. led trainees and contributed to colocalization analysis; I.M.H. led trainees and contributed to structural variant data; T.L. led trainees and provided feedback on manuscript; H.K.I. led trainees and led the GWAS analysis team; K.G.A. generated data, provided oversight of LDACC and pipelines, and provided feedback on manuscript; T.Q. helped with data interpretation and provided feedback on manuscript; K.K. helped with data interpretation and provided feedback on manuscript; S.B.M. co-led manuscript, led trainees, and co-wrote manuscript.

## Data and materials availability

All data used for these analyses are available through dbGaP (accession phs000424.v8) and the GTEx Portal (www.gtexportal.org).

