## Supplemental Materials for "Long non-coding RNA gene regulation and trait associations across human tissues"

### Materials and Methods

GTEx data

All human donors were deceased, with informed consent obtained via next-of-kin consent for the collection and banking of de-identified tissue samples for scientific research. The research protocol was reviewed by Chesapeake Research Review Inc., Roswell Park Cancer Institute's Office of Research Subject Protection, and the institutional review board of the University of Pennsylvania.

The v8 freeze of GTEx data includes whole genome sequencing (WGS) data from the whole blood of 838 post-mortem individuals, and RNA-sequencing (RNA-seq) data from 54 tissues. Each tissue has a different sample size for RNA-seq; we confined our analyses to the 49 tissues with N >70, for a total of 15,201 samples. For more details on donor characteristics, sample collection, and data production, see the GTEx v8 main paper (*18*).

For the tissue-specific expression analyses in this paper, the expression data used were gene-level TPM quantifications produced by RNA-SeQC (*44*) using the same GENCODE v26 collapsed single-transcript gene annotation from the GTEx v8 main paper (*18*), which are available on the GTEx Portal.

Defining gene groups

Five gene groups were compared throughout this paper: "total protein-coding genes", "expression-matched protein-coding genes", "total lncRNA genes", "high-confidence non-coding lncRNA genes", and "lncRNA genes with known function". The "total protein-coding genes" group includes any genes with the "protein_coding" biotype in the GTEx GENCODE v26 GTF (19,291 genes). The "total lncRNA genes" group includes any genes with a long non-coding gene biotype ("processed_transcript", "non_coding", "sense_intronic", "sense_overlapping", "antisense", "lincRNA", "macro_lncRNA", "bidirectional_promoter_lncRNA", "3prime_overlapping_ncRNA") in the GTEx GENCODE v26 GTF (14,100 genes).

The "expression-matched protein-coding genes" group were identified through the R package optmatch (*45*), which paired each expressed lncRNA with a protein-coding gene based on similarity of expression level (TPM). Expression matching was done separately in each tissue, which means that the same lncRNA could be matched with different protein-coding genes in different tissues. The union of expression-matched protein-coding genes across all 49 tissues was a set of 15,410 genes. Whenever possible, the tissue-specific sets of expression-matched protein-coding genes were used. The only time the entire union set is used at once is the tissue-specificity analyses of Figures 1B and 2D.

The "high-confidence non-coding lncRNA genes" come from LNCipedia's reference set of lncRNAs that have passed a series of particularly strict criteria to be defined as lacking protein-coding potential (4,510 genes) (*12*). The "lncRNA genes with known function" are a manually combined set of genes from lncRNAdb (*13*), the HUGO gene nomenclature committee (HGNC, https://www.genenames.org/data/genegroup/#!/group/788), and recent work that identified functional lncRNAs through splice-site-targeted CRISPR screens (*14*); plus 5 genes found in the literature that were not covered in these three sources (713 genes). Genes from lncRNAdb and the HGNC were only included if they had at least one reference in which direct manipulation of the gene (e.g., knockdown or overexpression) had some effect on cellular phenotype or other genes' expression.

Identifying independent *cis*-eQTLs by forward stepwise regression-backward selection

The same independent cis-eQTLs mapped for the main GTEx paper were used in this paper; for full details of the methods used, see the GTEx v8 main paper (*18*). In each tissue, the variants tested were those within 1 Mb of the TSS of each gene and with minor allele frequencies ≥0.01, with the minor allele observed in at least 10 samples of that tissue.

Independent eQTLs were used for Figure 2B,C, as well as the eVariant-sharing analyses of Figures 4 and S5, and Table S5. eVariant sharing was defined as one gene's associated eVariant either being 1) an eVariant for another gene in the same tissue, or 2) being linked (variants within 500kb of each other and an R2 ≥0.85) to an eVariant for another gene within the same tissue.

Identifying tissue-shared *cis*-eQTLs with mash

To identify tissue-specific and tissue-shared eQTLs, multivariate adaptive shrinkage (mash) was used (*46*). This flexible method is designed to deal with large numbers of tests that span many different conditions. In this case, it was used to produce estimates of both effect size and significance of variant-gene associations across the different tissue conditions.

The mash results from the main GTEx paper were used in this paper; for full details of the methods used, see the GTEx v8 main paper (*18*). The output of mash is local false sign rate (LFSR), which is analogous to the false discovery rate, as well as a beta-value effect size. Variant-gene associations with LFSR ≤0.05 were considered significant. The mash output were used in analyses of tissue specificity of eQTLs - this includes Figures 2A,D,E, Figure S3, and Table S3.

Multi-tissue gene expression outlier discovery

We subset expression data in each tissue to genes with 6 reads and TPM > 0.1 in at least 10% of individuals. Within each tissue, the TPM values were log transformed, (log2(TPM + 2)), and scaled across individuals for each gene. We regressed out the effects of the first three genotype principal components, and hidden factors discovered via PEER (*47*), the number of which depends on sample size for that tissue and was consistent with the GTEx eQTL discovery pipeline (*18*): for tissues with less than 150 samples, we removed 15 PEER factors; less than 250 samples, 30 factors; less than 350, 45 factors; and 60 for the remaining tissues. We additionally corrected for the genotype of the strongest cis-eQTL per gene per tissue to magnify rare variant effects, which has been shown to improve nearby rare variant enrichments (*22*). We then re-scaled expression values across individuals within each gene to generate corrected Z-scores per individual per gene per tissue.

For each gene-individual pair, if that individual has expression measurements in at least five tissues, we calculate a median Z-score for that gene. We define outlier individuals as those with a |median Z-score| greater than 2, and non-outliers as all other individuals for the same set of genes. We removed as global outliers 39 individuals for whom the proportion of tested genes were outliers at a threshold of |median Z-score| > 3 exceeded 1.5 times the interquartile range of the distribution of proportion outlier genes per individual. For outlier analysis, we include all autosomal lincRNA and protein-coding genes.

To assess variant enrichments, we subset to 714 individuals who self-report with European ancestry, as allele frequencies are less comparable between continental populations. We retain all SNPs and indels that pass quality control in the GTEx VCF. Structural variants were called in a subset of these individuals as in (*48*) with GenomeSTRiP GSCNQUAL (*49*) set to limit the false discovery rate (FDR) for each variant type. GenomeSTRiP's IntensityRankSumAnnotator was used to evaluate FDR based on available Illumna Human Omni 5M gene expression array data. GSCNQUAL was limited to ≥ 1 for GenomeSTRiP deletions and ≥ 8 for multi-allelic copy number variants, corresponding to an FDR of 10%. The GSCNQUAL cutoff for GenomeSTRiP duplications was set at ≥ 17, the point where the FDR plateaued at 15.1% and did not fluctuate more than ±1% for over 50 steps in increasing GSCNQUAL score. Additionally, the Mobile Element Locator Tool (MELT) (*50*) version 2.1.4 was run using MELT-SPLIT to identify ALU, SVA, and LINE1 insertions into the test genomes. MELT calls that were categorized as "PASS" in the VCF info field, had an ASSESS score ≥ 3, and SR count ≥ 3 were retained.

We define rare variants as those with < 1% frequency in GTEx, and for SNPs and indels, also < 1% frequency in non-Finnish Europeans from the gnomAD database (*51*). Remaining bins are defined by GTEx allele frequencies. We calculate relative risk as the proportion of outlier individuals with a variant of a given frequency within 10kb of the outlier gene or in the gene body over the proportion of non-outlier individuals with a variant of a given frequency within 10kb of or in the gene body of the same set of genes. Variant categories were annotated using Ensembl VEP (version 88) and each gene-individual pair was assigned to the most enriched variant category, regardless of the number of nearby rare variants.

Allele-specific expression (ASE)

Autosomal ASE data were produced using GATK ASEReadCounter tool (*52*) and the WASP filtering strategy (*53*) to remove read mapping bias, as described in the GTEx main paper (*18*) and the GTEx ASE companion paper (*54*). ASE sites were removed if they were in low-mappability regions (75-mer mappability with leq2 mismatches < 1), showed mapping bias in simulation (*55*), or had no more reads supporting two alleles than would be expecting from sequencing noise alone, indicating potential genotyping error (FDR < 1%, see (*52*) for description of test). Phase was not incorporated in these ASE analyses.

To evaluate maintenance of ASE along neighboring genes, 'central genes' were identified in each sample (individual-tissue combination) that had strong ASE (multiple test-adjusted binomial p-value ≤0.05 and allele ratio either 0.02-0.15 or 0.85-0.98 for any variant in the gene). For each central gene, all non-overlapping genes with a TSS within 500 kb of the central gene's TSS were identified as neighboring genes. Neighbor genes with ASE data available in that sample were then checked for the presence of any variant with significant ASE (multiple test-adjusted binomial p-value ≤0.05) to calculate the proportion of neighboring genes with significant ASE.

Note that since ASE is an individual-specific measurement, the central genes selected and the ASE data available for neighboring genes varied for each sample, and the ASE plots are based on aggregates across all samples.

Weighted gene co-expression network analysis (WGCNA)

Transcript per million (TPM) values quantified by RNASeQC (*44*) were normalized on a per tissue basis using the variance-stabilized normalization (VSN) as implemented by the vsn package (*56*). Only genes which met an expression cutoff of at least 0.1 TPM in at least 20% of samples were included. The effects of gene expression batch, Hardy death type, and ischemic time were removed from normalized TPM values in each tissue using an empirical Bayes linear model implemented by the *WGCNA* package (*28*). Latent factors were not removed from the expression data, because we found that doing so eliminated biological signals necessary for constructing the co-expression networks.

Adjacency matrices were computed using biweight mid-correlation and the default soft-thresholding power of 12. The adjacency matrix was transformed into a topological overlap matrix (TOM) and then subtracted from 1 to create a dissimilarity TOM suitable for hierarchical clustering.

Co-expression modules were identified using the dynamic tree-cutting approach provided by the WGCNA package (*28, 57*). The dissimilarity TOM was transformed into a Euclidean distance matrix and a hierarchical clustering tree was created from this matrix using average-linked hierarchical clustering. The hybrid dynamic tree-cutting algorithm was used with a minimum module size of 50 to prevent the creation of very small modules and a deepSplit parameter of 3 to favor more small modules over few large modules. The pamRespectsDendro parameter was set to true, which will force the partitioning around medoids (PAM) step to respect the hierarchical clustering tree when attempting to assign unclustered genes to modules or to merge very similar modules. This is more conservative than setting pamRespectsDendro to false and leads to more genes remaining unclustered.

Eigengenes were computed from the first principal component of the expression values of the genes assigned to each module. Modules whose eigengenes had a biweight mid-correlation greater than 0.8 were merged using the mergeCloseModules function to reduce the number of highly correlated modules. Module membership was estimated as the biweight midcorrelation between each gene and its module eigengene. Scaled intramodular connectivity ($k_{in}$) was computed from intramodular connectivity for each gene by dividing its intramodular connectivity by the largest intramodular connectivity value in that module (scaled $k_{in}$ range = 0 to 1).

<u>Co-expression network module annotation via gene set enrichment</u>

In large modules, gene set enrichment was limited to the top 500 genes as ranked by module membership. Enrichments for both cell compartments and cell types were assessed.

For cell compartment gene sets, only five terms were tested: nucleolus (GO:0005730), mitochondrion (GO:0005739), mitochondrial inner membrane (GO:0005743), and cytosolic ribosome (GO:0022626). This was because our aim was to eliminate spurious cell type enrichment driven by cell compartment rather than cell types. Enrichment was computed using the hypergeometric overlap test between the genes in the module and list of cell compartment genes provided by GO Cellular Compartment (*58, 59*). Enrichment p-values were adjusted for the number of modules and the number of terms in each tissue network using Bonferroni correction.

Enrichment for cell types across 20 tissues were computed using cell type specificity index (SI) values estimated by the pSI package (*60, 61*) in the datasets described in the Supplementary Methods ('Cell type annotation sources'). For each module, a linear model was fitted with a dummy variable indicating membership in the module or (or the top 500 genes in the module for large modules) as the predictor, and the SI values as the outcome. Only genes found in both the network and annotation were used in the model. Because a lower SI value indicates a higher cell type specificity, only models with a negative coefficient were considered a valid enrichment. All enrichment p-values were Bonferroni-corrected for the number of modules and the number of cell types in each dataset.

Final annotations were decided by combining the cell compartment and cell type annotations. Modules with strong enrichment for mitochondria or ribosomes were annotated for those compartments over cell types. Cell types present in multiple tissues from the annotation sources (such as resident immune cells, epithelial cells, or stromal cells) were accepted as annotations if there was agreement from multiple tissues. Tissue-specific cell types were only used as a module annotation if they were in the appropriate tissue. For tissues without a direct analog in the annotation sources, cell type annotations were assigned by looking across all available tissues in the annotation sources. If a module was annotated for a mix of cell types, it was submitted to Enrichr (*62, 63*) to assist in identifying other pathways. Cell types or pathways identified indirectly were annotated as "uncertain" to reflect the lower confidence in their annotation.

Cell type annotation sources

Blood cell type-specific expression data was obtained from a published dataset (GSE24759) (*64*). Raw CEL files were imported and normalized with RMA using the affy package (*65*). Progenitor cells and cell types with a small number of samples were excluded, and some cell subtypes were aggregated. The final dataset used to estimate specificity index values was the averaged expression values for the samples corresponding to the following 13 cell types: naive CD4+ T-cell, memory CD4+ T-cell, naive CD8+ T-cell, memory CD8+ T-cell, naive B-cell, mature B-cell, mature NK cell, monocyte, myeloid dendritic cell, granulocyte (neutrophil), basophil, eosinophil, and megakaryocyte.

Cell type-specific expression for the central nervous system was obtained from a published single-cell RNA dataset (*66*). Fetal cell types were excluded and log counts were averaged for estimating specificity index values for six cell types: neuron, astrocyte, oligodendrocyte, oligodendrocyte progenitor cell (OPC), microglia and endothelial cell.

The remaining cell type specific expression data was obtained from the Mouse Cell Atlas (*28*). Counts data was downloaded from Mouse Cell Atlas website (http://bis.zju.edu.cn/MCA/) and log normalized. All adult tissues with a matching tissue in GTEx were used except peripheral blood and brain and neonatal heart was also included because no adult heart sample was available. Some cell subtypes were collapsed, and averaged cell type matrices were computed for 18 tissues: heart, kidney, liver, lung, mammary gland (involution), mammary gland (lactation), mammary gland (pregnancy), mammary gland (virgin), muscle, ovary, skin, pancreas, prostate, small intestine, spleen, stomach, testis, and uterus.

Colocalization of GWAS and eQTL signals

Colocalization was performed between eQTL and GWAS signals across 87 traits and 49 tissues. The colocalization results from the main GTEx paper were used in this paper; for full details of the methods and a complete list of the GWAS studies used, see the GTEx v8 main paper (*19*) and GWAS companion paper (*33*). We used a threshold of posterior probability of colocalization (PP4) ≥0.5 to define significant colocalization events.

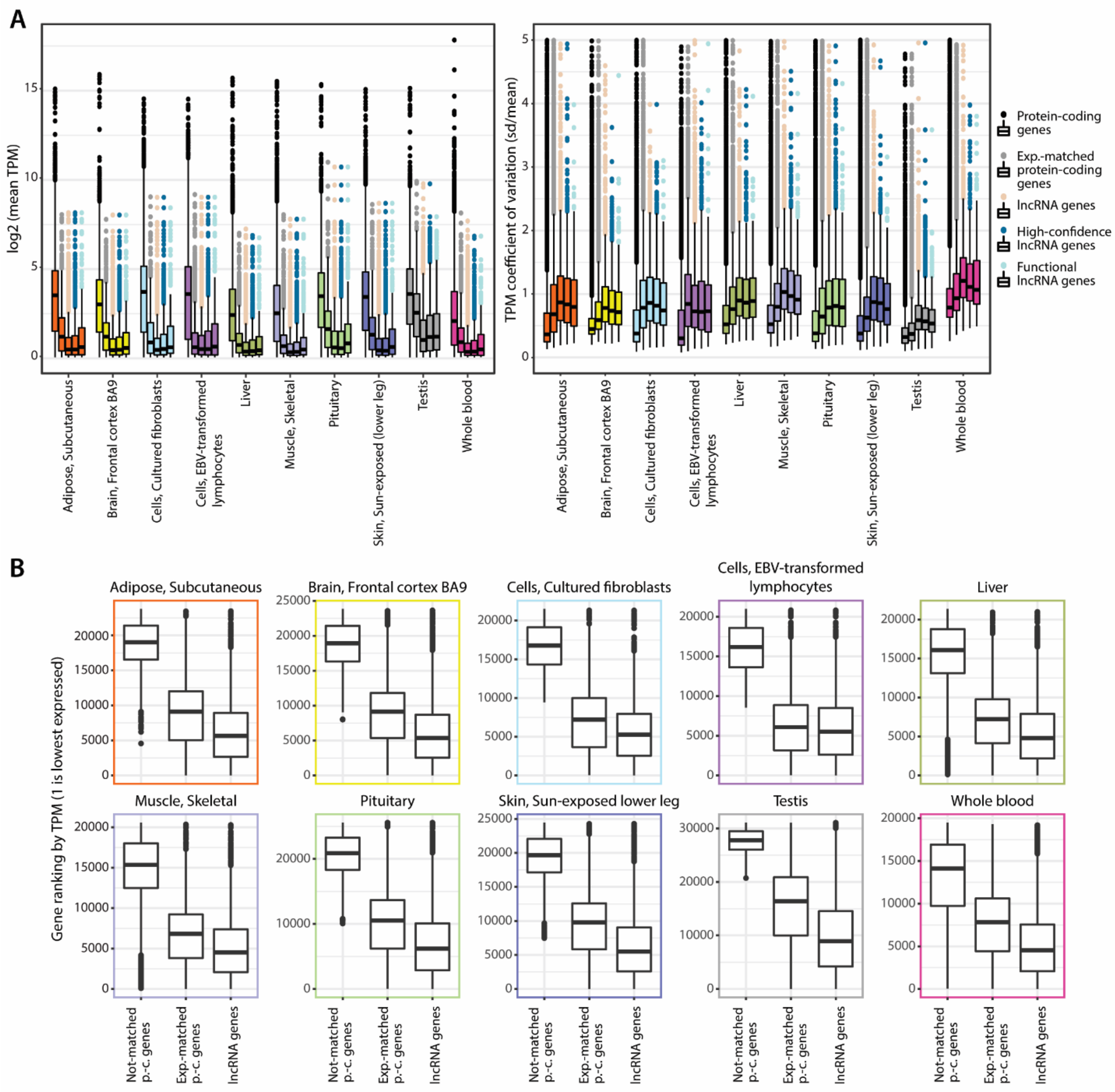


**Fig. S1.**

Creating expression-matched sets of protein-coding genes. (A) Comparison of mean gene expression levels *(left)* and gene expression variation *(right)* across gene groups in a subset of GTEx tissues. The y-axis of the coefficient of variation plot is truncated at 5 for visibility; the maximum value is 25.5. (B) Evaluating the expression-matched protein-coding genes. Within each tissue, all protein-coding genes and lncRNAs were ranked by mean expression (TPM), with a ranking of 1 assigned to the gene with the lowest mean expression. For each tissue depicted, the box plots show the rankings of (from left to right): protein-coding genes that did not match with any expressed lncRNA in that tissue; the protein-coding genes that matched with an expressed lncRNA; and the expressed lncRNA genes. p.-c. genes = protein-coding genes.

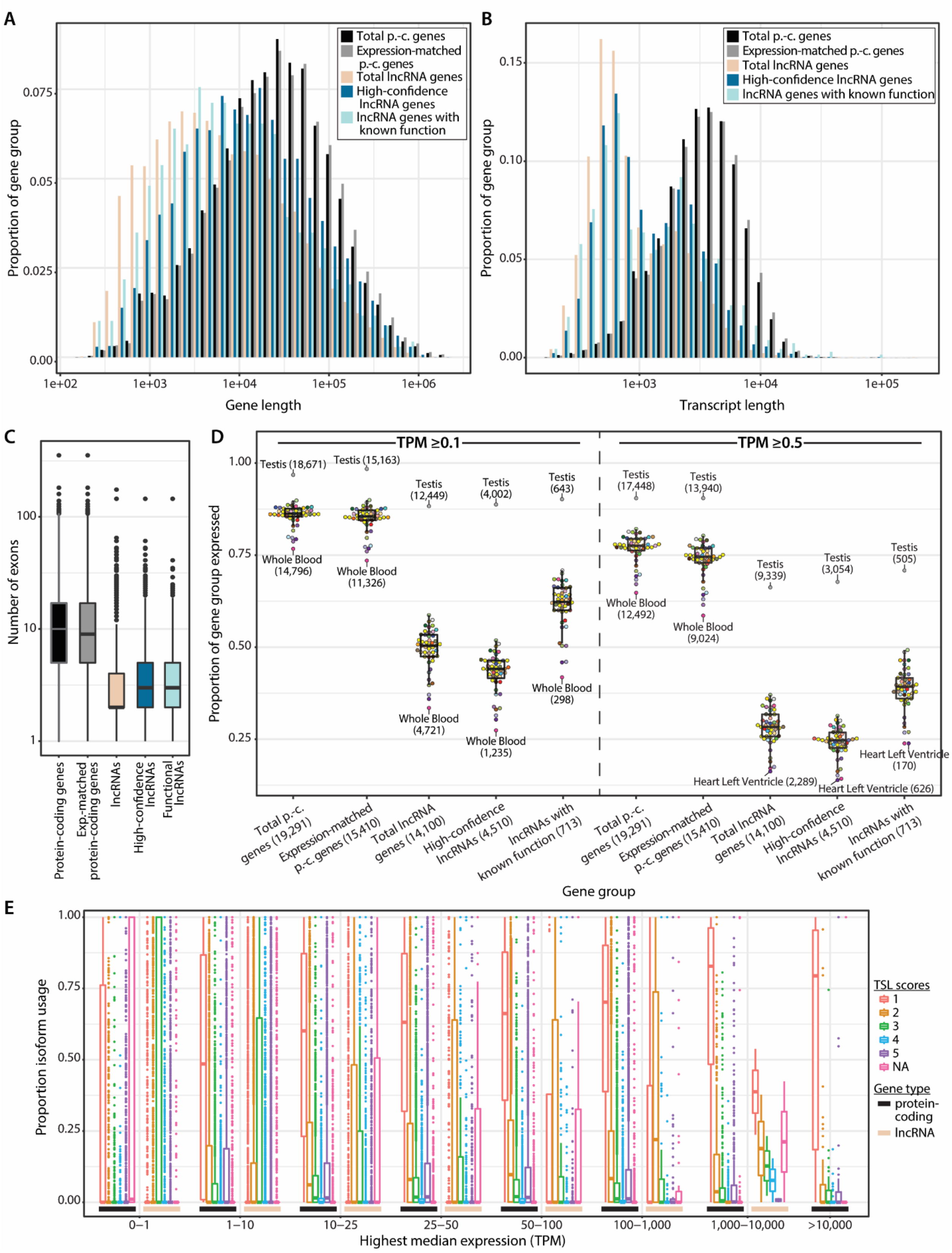
A
Total p.-c. genes
Expression-matched p.-c. genes
Total lncRNA genes
High-confidence lncRNA genes
lncRNA genes with known function
Proportion of gene group
Gene length
B
Total p.-c. genes
Expression-matched p.-c. genes
Total lncRNA genes
High-confidence lncRNA genes
lncRNA genes with known function
Proportion of gene group
Transcript length
C
Number of exons
Protein-coding genes
Exp.-matched protein-coding genes
lncRNAs
High-confidence lncRNAs
Functional lncRNAs
D
TPM ≥0.1
TPM ≥0.5
Proportion of gene group expressed
Testis (18,671)
Testis (15,163)
Testis (12,449)
Testis (4,002)
Testis (643)
Testis (17,448)
Testis (13,940)
Testis (9,339)
Testis (3,054)
Testis (505)
Whole Blood (14,796)
Whole Blood (11,326)
Whole Blood (4,721)
Whole Blood (1,235)
Whole Blood (298)
Whole Blood (12,492)
Whole Blood (9,024)
Heart Left Ventricle (2,289)
Heart Left Ventricle (626)
Heart Left Ventricle (170)
Total p.-c. genes (19,291)
Expression-matched p.-c. genes (15,410)
Total lncRNA genes (14,100)
High-confidence lncRNAs (4,510)
lncRNAs with known function (713)
Gene group
E
Proportion isoform usage
TSL scores
1
2
3
4
5
NA
Gene type
protein-coding
lncRNA
0–1
1–10
10–25
25–50
50–100
100–1,000
1,000–10,000
>10,000
Highest median expression (TPM)

**Fig. S2.**

Characteristics of lncRNA genes and protein-coding genes. Distributions of (A) gene length (genomic start position to genomic end position) and (B) transcript length, by gene group. (C) Range in number of exons across all genes in a given gene group. (D) Proportion of genes expressed in each gene group. Data points reflect the proportion of a gene group expressed in a given tissue, with point color indicating the tissue. The overlaying box plot summarizes the values across tissues. The most extreme values are labeled with the tissue name and number of genes expressed. (E) Distribution of transcript support level (TSL) scores across protein-coding and lncRNA gene transcripts, binned by the gene's highest median expression across all tissues. For each gene, the proportion of expressed isoforms were calculated across all tissues, the isoforms' TSL scores were obtained from Ensembl Archive 88, and proportions were summed across isoforms with the same TSL score. p.-c. genes = protein-coding genes.

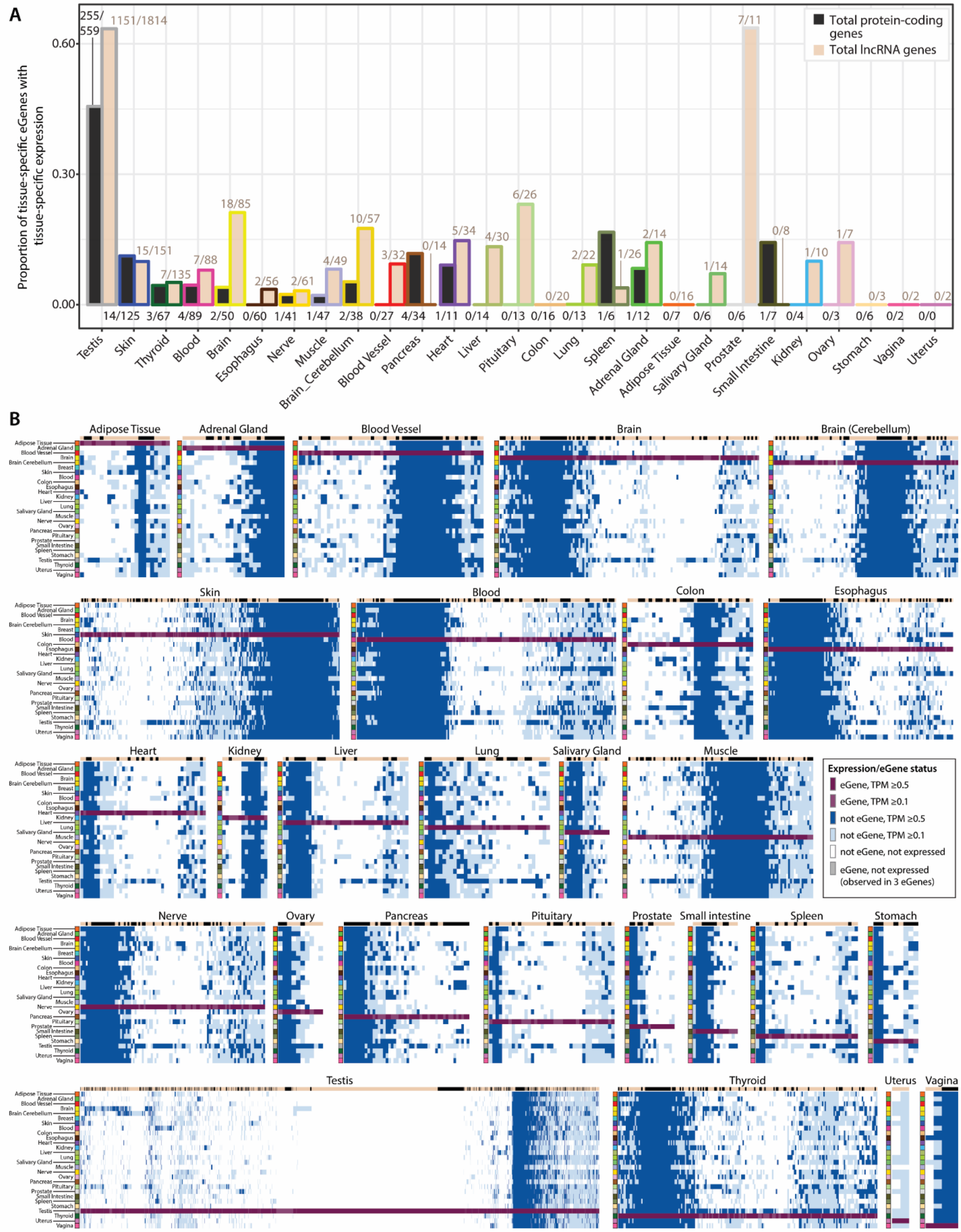
A
Proportion of tissue-specific eGenes with tissue-specific expression
0.60
0.30
0.00
Total protein-coding genes
Total lncRNA genes
255/559
1151/1814
15/151
7/135
7/88
18/85
2/56
2/61
4/49
10/57
3/32
0/14
5/34
4/30
6/26
0/20
2/22
1/26
2/14
0/16
1/14
7/11
0/8
1/10
1/7
0/3
0/2
0/2
14/125 3/67 4/89 2/50 0/60 1/41 1/47 2/38 0/27 4/34 1/11 0/14 0/13 0/16 0/13 1/6 1/12 0/7 0/6 0/6 1/7 0/4 0/3 0/6 0/2 0/0
Testis Skin Thyroid Blood Brain Esophagus Nerve Muscle Brain_Cerebellum Blood Vessel Pancreas Heart Liver Pituitary Colon Lung Spleen Adrenal Gland Adipose Tissue Salivary Gland Prostate Small Intestine Kidney Ovary Stomach Vagina Uterus
B
Adipose Tissue
Adrenal Gland
Blood Vessel
Brain
Brain (Cerebellum)
Skin
Blood
Colon
Esophagus
Heart
Kidney
Liver
Lung
Salivary Gland
Muscle
Nerve
Ovary
Pancreas
Pituitary
Prostate
Small intestine
Spleen
Stomach
Testis
Thyroid
Uterus
Vagina
Expression/eGene status
eGene, TPM ≥0.5
eGene, TPM ≥0.1
not eGene, TPM ≥0.5
not eGene, TPM ≥0.1
not eGene, not expressed
eGene, not expressed (observed in 3 eGenes)

**Fig. S3.**

Broad tissue-specific eGenes that also show broad tissue-specific expression. (A) Proportion of the broad tissue-specific eGenes (from Fig. 2E) that also show broad tissue-specific gene expression (displayed in Fig. 1C). Tissues are ordered from those with the highest number of tissue-specific eGenes (testis) to those with the lowest (uterus). Expression threshold was $\geq$0.5 TPM in >20% of samples. Labels show (number of broad tissue-specific eGenes that also have broad tissue-specific expression) / (total number of broad tissue-specific eGenes). (B) Expression of each broad tissue's tissue-specific eGenes across all tissues. For each tissue's heatmap, the columns represent eGenes were only discovered in that broad tissue.

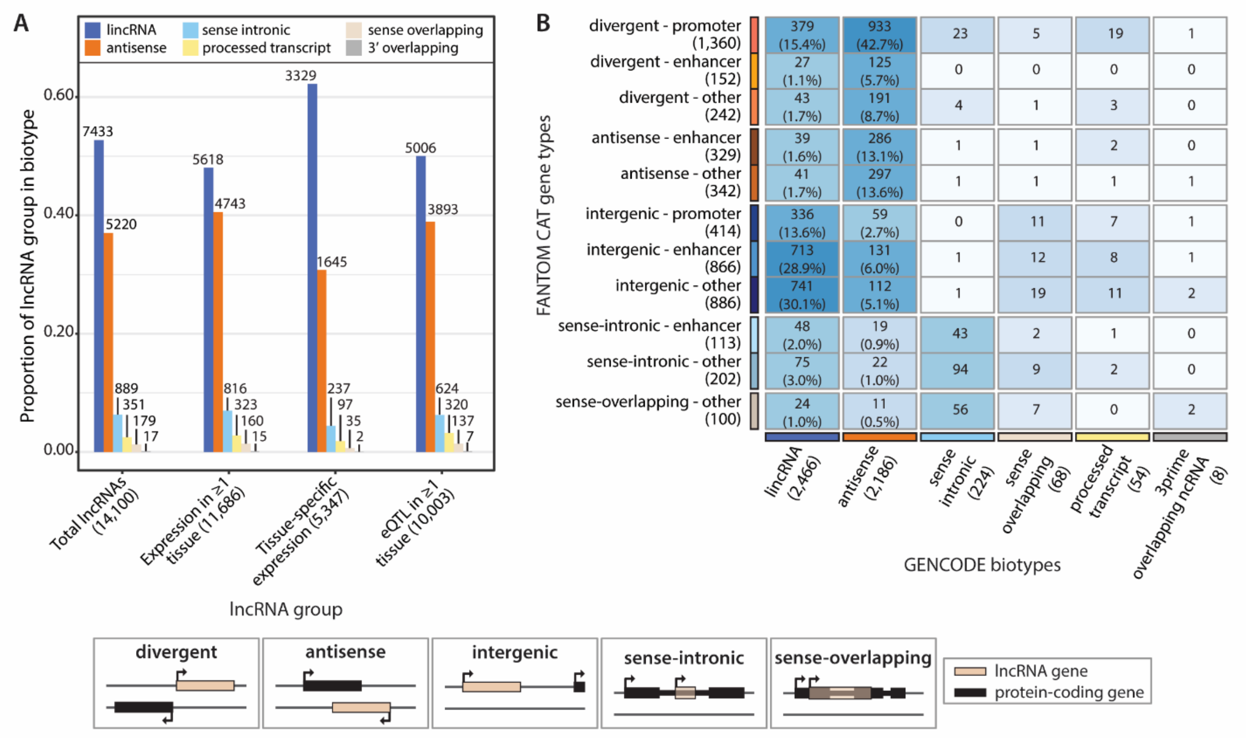


| FANTOM CAT gene types | lincRNA (2,466) | antisense (2,186) | sense intronic (224) | sense overlapping (68) | processed transcript (54) | 3prime overlapping ncRNA (8) |
|---|---|---|---|---|---|---|
| divergent - promoter (1,360) | 379 (15.4%) | 933 (42.7%) | 23 | 5 | 19 | 1 |
| divergent - enhancer (152) | 27 (1.1%) | 125 (5.7%) | 0 | 0 | 0 | 0 |
| divergent - other (242) | 43 (1.7%) | 191 (8.7%) | 4 | 1 | 3 | 0 |
| antisense - enhancer (329) | 39 (1.6%) | 286 (13.1%) | 1 | 1 | 2 | 0 |
| antisense - other (342) | 41 (1.7%) | 297 (13.6%) | 1 | 1 | 1 | 1 |
| intergenic - promoter (414) | 336 (13.6%) | 59 (2.7%) | 0 | 11 | 7 | 1 |
| intergenic - enhancer (866) | 713 (28.9%) | 131 (6.0%) | 1 | 12 | 8 | 1 |
| intergenic - other (886) | 741 (30.1%) | 112 (5.1%) | 1 | 19 | 11 | 2 |
| sense-intronic - enhancer (113) | 48 (2.0%) | 19 (0.9%) | 43 | 2 | 1 | 0 |
| sense-intronic - other (202) | 75 (3.0%) | 22 (1.0%) | 94 | 9 | 2 | 0 |
| sense-overlapping - other (100) | 24 (1.0%) | 11 (0.5%) | 56 | 7 | 0 | 2 |

**Fig. S4.**

Comparison of GENCODE and FANTOM CAT lncRNA gene types. (A) Proportion of gene “hit lists” made up of each lncRNA GENCODE subtype. Bar labels indicate the number of lncRNA genes. (B) Comparison of lncRNA subtype classifications in GENCODE (used in GTEx) versus FANTOM CAT (*1*). Only lncRNAs with subtype assignments in both annotations are included. For the GENCODE subtypes of lincRNA and antisense, the percentage of the subtype in each FANTOM CAT category is also labeled.

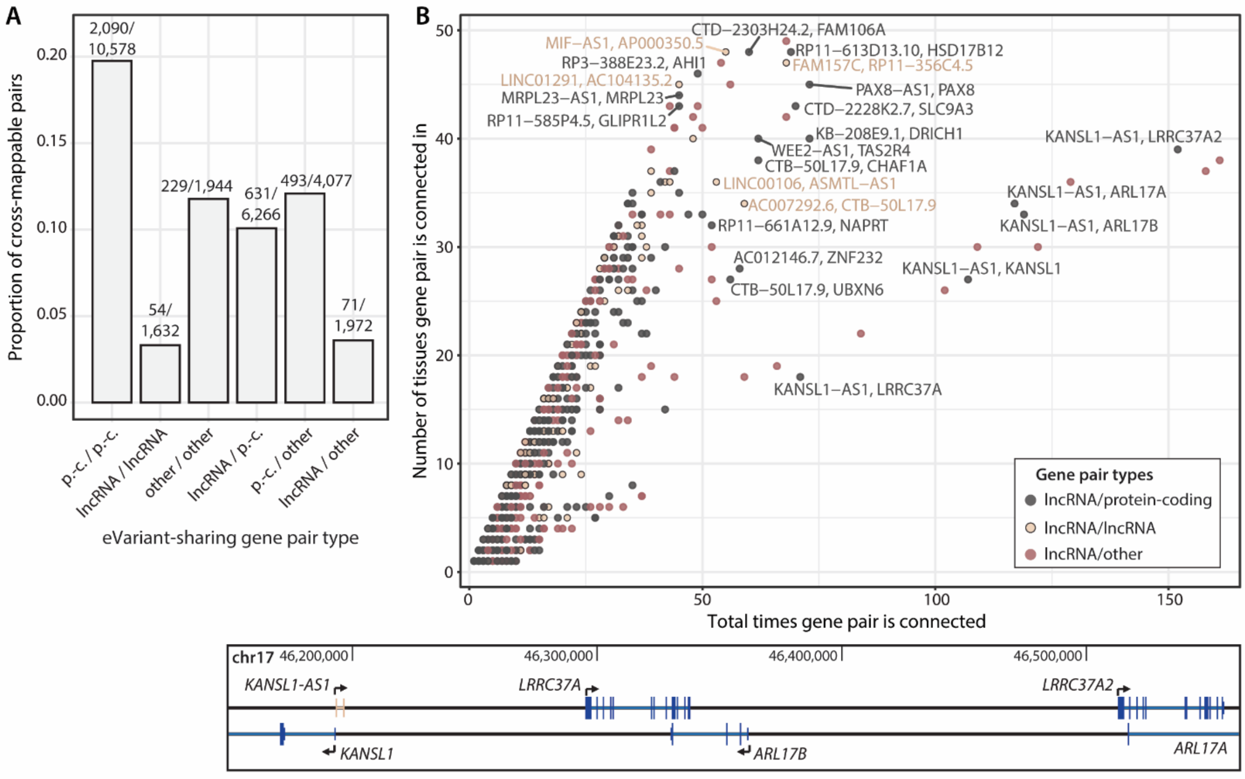


**Fig. S5.**

Frequency of eVariant sharing. (A) Proportion of eVariant-sharing gene pairs that are cross-mappable (symmetric cross-mappability score >0), grouped by the gene pair type. Labels indicate the number of cross-mappable gene pairs / total number of gene pairs. (B) *(top)* Recurring gene pairs that share an eVariant and involve at least one lncRNA gene. Since a gene pair can share more than 1 eVariant in the same tissue, sometimes the total times a gene pair is connected (x-axis value) exceeds the number of tissues the gene pair is connected in (y-axis value). lncRNA/lncRNA and lncRNA/protein-coding gene pairs that occur in >40 tissues, or that occur >50 times total, are labeled. *(bottom)* Genomic locations of a group of genes on chromosome 17 (including one lncRNA, *KANSL1-AS1*) that frequently shared eVariants. p.-c. genes = protein-coding genes.

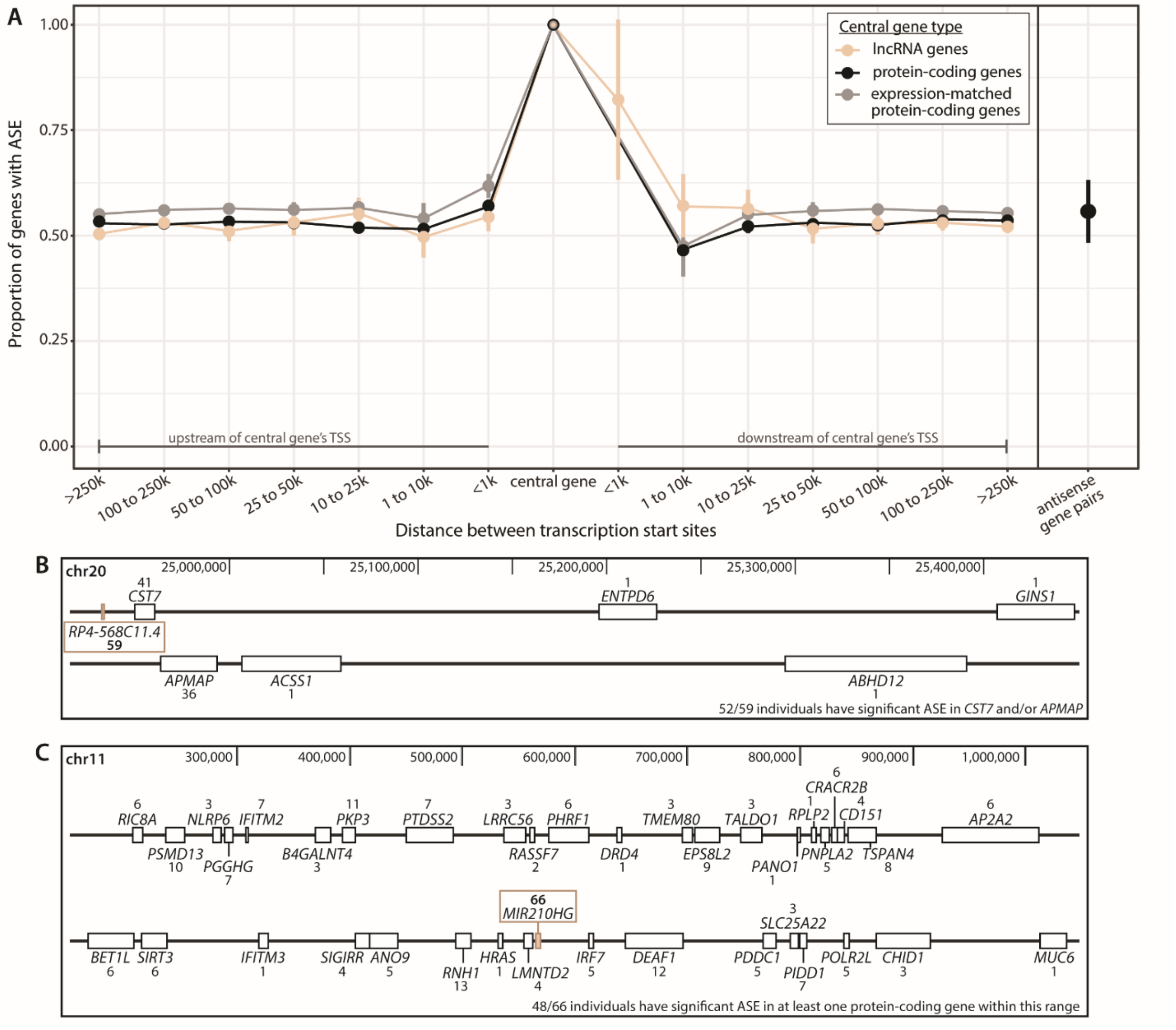


**Fig. S6.**

Extension of allele-specific expression (ASE) from protein-coding and lncRNA genes. Central genes were identified as either protein-coding or lncRNA genes containing a variant with significant strong ASE (adjusted binomial p-value ≤0.05, allelic ratio deviation (from 0.5) of 0.35-0.48). (A) Proportion of non-overlapping neighboring protein-coding genes with significant ASE, binned by distance of the neighbor gene's TSS from the central gene's TSS. Coloring indicates the central gene's type (lncRNA, protein-coding, or expression-matched protein-coding). The right panel is specific to antisense-sense gene relationships: the central genes are antisense lncRNA genes with strong ASE, and only the sense protein-coding gene neighbors were used in ASE calculation. The bars indicate the 95% CI across different central genes, with ASE status being collapsed by individuals and tissues. (B) A frequently co-occurring set of genes with ASE, which was identified starting from the lncRNA central gene *RP4-568C11.4*. Only genes tested for ASE in at least one individual are shown. Numbers indicate the number of individuals that showed significant ASE for the labeled gene. (C) A region of long-range ASE containing a lncRNA gene with significant strong ASE that was observed in multiple individuals. *MIR210HG* was a lncRNA central gene in 66 individuals; of these, 48 had significant ASE in at least one protein-coding gene within the range depicted. Only genes with significant ASE in at least one individual are shown. Numbers indicate the number of individuals that showed significant ASE for the labeled gene. For (B) and (C), the lncRNA central gene is colored brown.

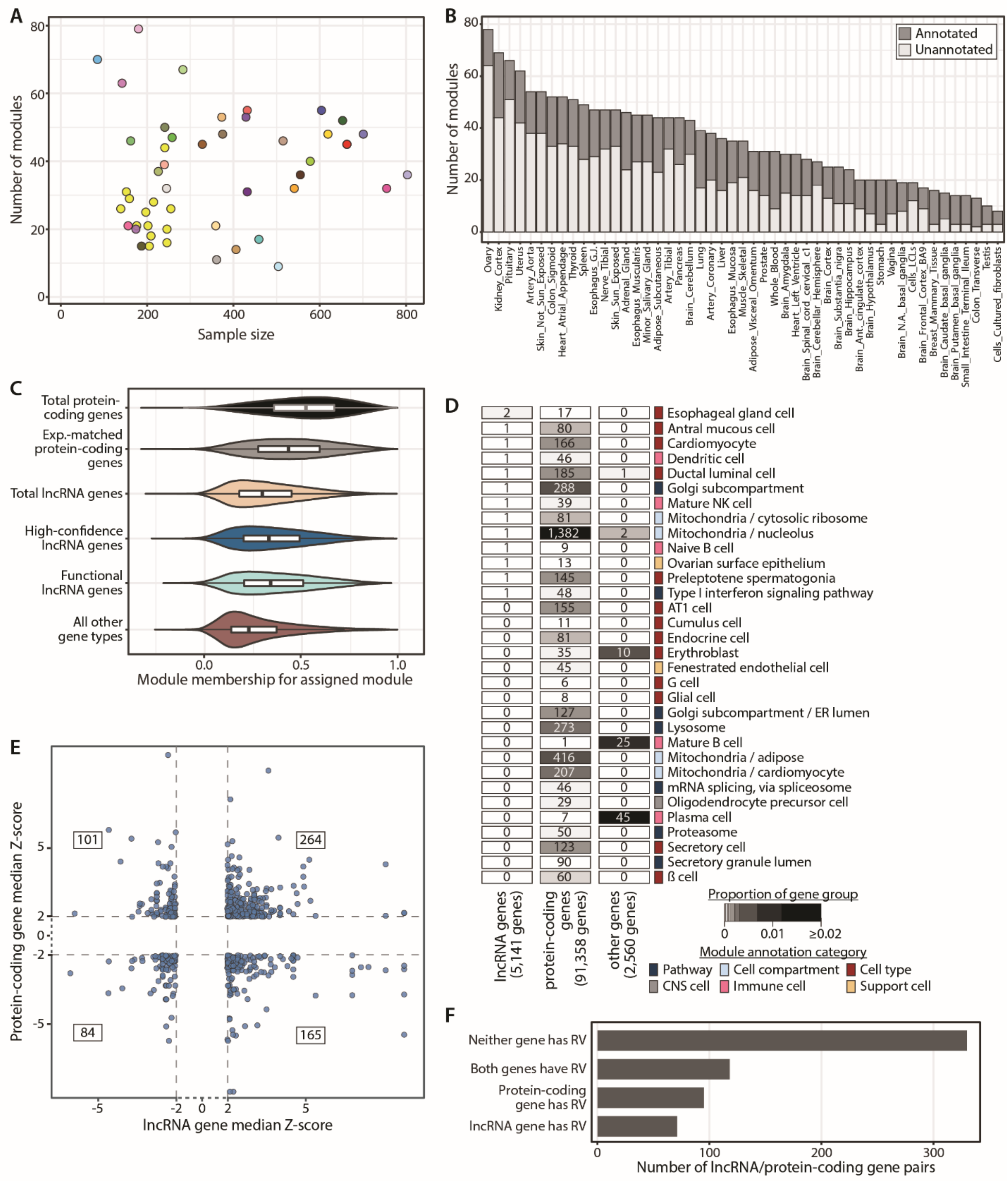


**Fig. S7.**

Additional figures from weighted gene co-expression network analysis (WGCNA). (A) The number of co-expression modules identified in each tissue is not related to the sample size of the tissue. Fill color of points indicates tissue. (B) Number of modules with a cell type or cell compartment annotation versus those that are unannotated in each tissue. There is also one module of unclustered genes per tissue, which is not depicted. (C) Distribution of module membership scores of each gene for its assigned module, separated by gene group. (D) Continuation of Figure 5D: module annotations of genes with high intra-modular connectivity (the gene is in the top

$k_{in}$ rank decile of its module, and has scaled $k_{in}$ ≥0.5). Box fill reflects the proportion of genes assigned to a module with that annotation. Since genes are assigned to modules in multiple tissues, the labels reflect gene-tissue combinations, not individual genes. (E) Proportion of gene pairs assigned to the same module based on whether they share an eVariant. The control gene pairs are matched such that their first gene is from one of the gene pairs that shares an eVariant, and the second gene is within 500kb of the first gene and has an eQTL. The “known antisense pairs” type is a set of established protein-coding/antisense-lncRNA gene pairs. (F) Median Z-scores of the 614 lncRNA/protein-coding gene pairs that are outliers and share >12 co-expression modules. (G) Presence of rare variants (RV) in the 614 outlier events involving lncRNA/protein-coding gene pairs that share >12 modules. Just over half of the individuals do not have an RV near either gene, and the rest have an RV near one or both genes.

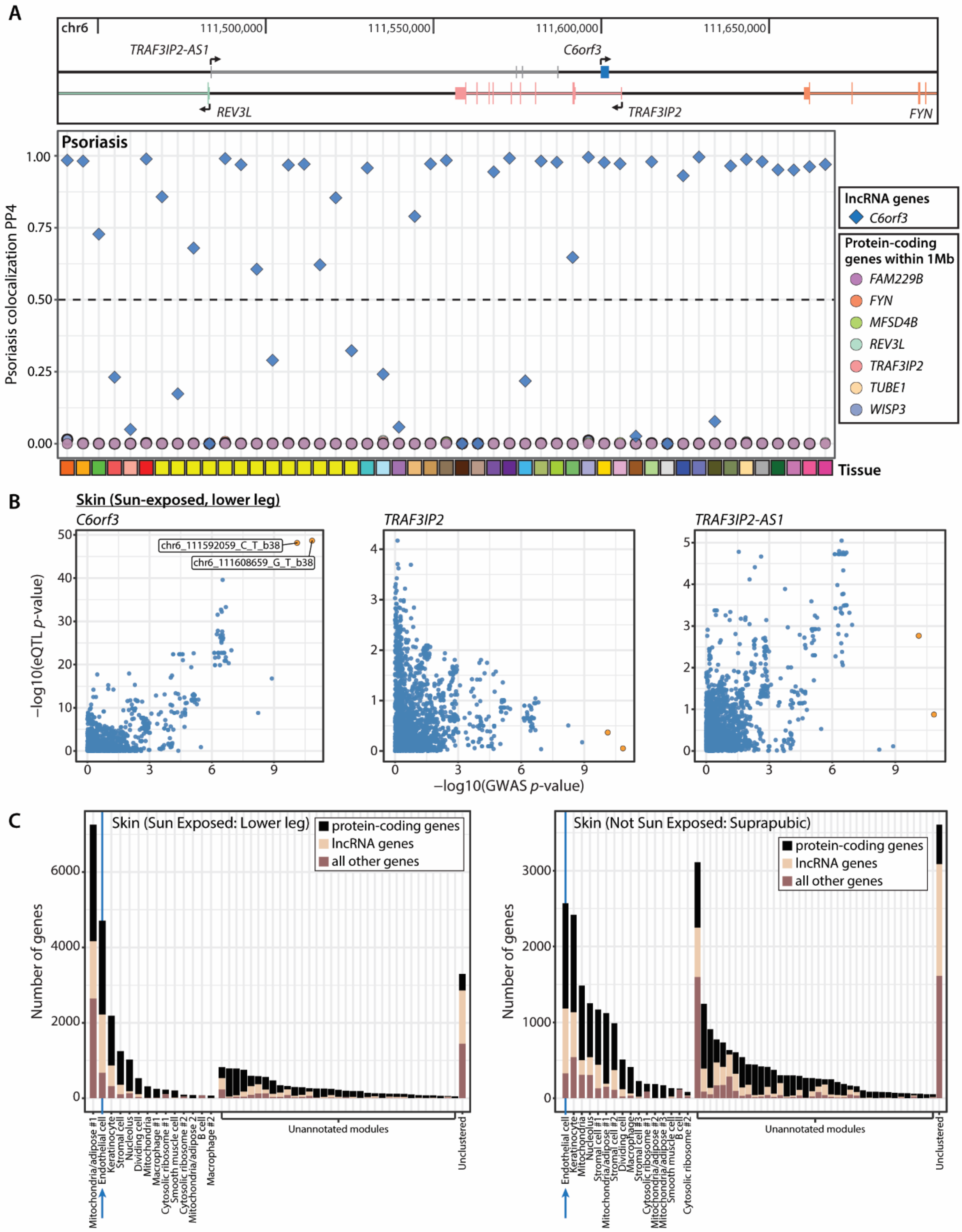
A
chr6
111,500,000
111,550,000
111,600,000
111,650,000
TRAF3IP2-AS1
C6orf3
REV3L
TRAF3IP2
FYN
Psoriasis
Psoriasis colocalization PP4
lncRNA genes
C6orf3
Protein-coding genes within 1Mb
FAM229B
FYN
MFSD4B
REV3L
TRAF3IP2
TUBE1
WISP3
Tissue
B
Skin (Sun-exposed, lower leg)
C6orf3
TRAF3IP2
TRAF3IP2-AS1
chr6_111592059_C_T_b38
chr6_111608659_G_T_b38
−log10(eQTL p-value)
−log10(GWAS p-value)
C
Skin (Sun Exposed: Lower leg)
Skin (Not Sun Exposed: Suprapubic)
protein-coding genes
lncRNA genes
all other genes
Number of genes
Unannotated modules
Unclustered
Mitochondria/adipose #1
Endothelial cell
Keratinocyte
Stromal cell
Nucleolus
Dividing cell
Mitochondria
Macrophage #1
Cytosolic ribosome #1
Smooth muscle cell
Cytosolic ribosome #2
Mitochondria/adipose #2
B cell
Macrophage #2
Stromal cell #1
Stromal cell #2
Macrophage
Stromal cell #3
Mitochondria/adipose #3

**Fig. S8.**

*C6orf3* is a single-exon lncRNA gene with significant colocalization events for psoriasis. (A) *(top)* Location of *C6orf3* and nearby protein-coding genes. *(bottom)* Colocalization posterior probability values across each tissue for *C6orf3,* as well as all protein-coding genes within 1Mb of the lncRNA gene, with psoriasis. The dashed line indicates the threshold for significance, PP4 $\leq 0.5$. (B) Plots of eQTL significance (in the sun-exposed skin tissue) versus psoriasis-GWAS significance for *C6orf3*, its overlapping protein-coding gene *TRAF3IP2*, and the nearby antisense lncRNA gene *TRAF3IP2-AS1*. The two most significant GWAS variants are colored in red. (C) WGCNA modules of the two skin tissues, ordered by annotation and size. The blue arrows indicate the modules that *C6orf3* was assigned to - an "endothelial cell" module in both tissues.

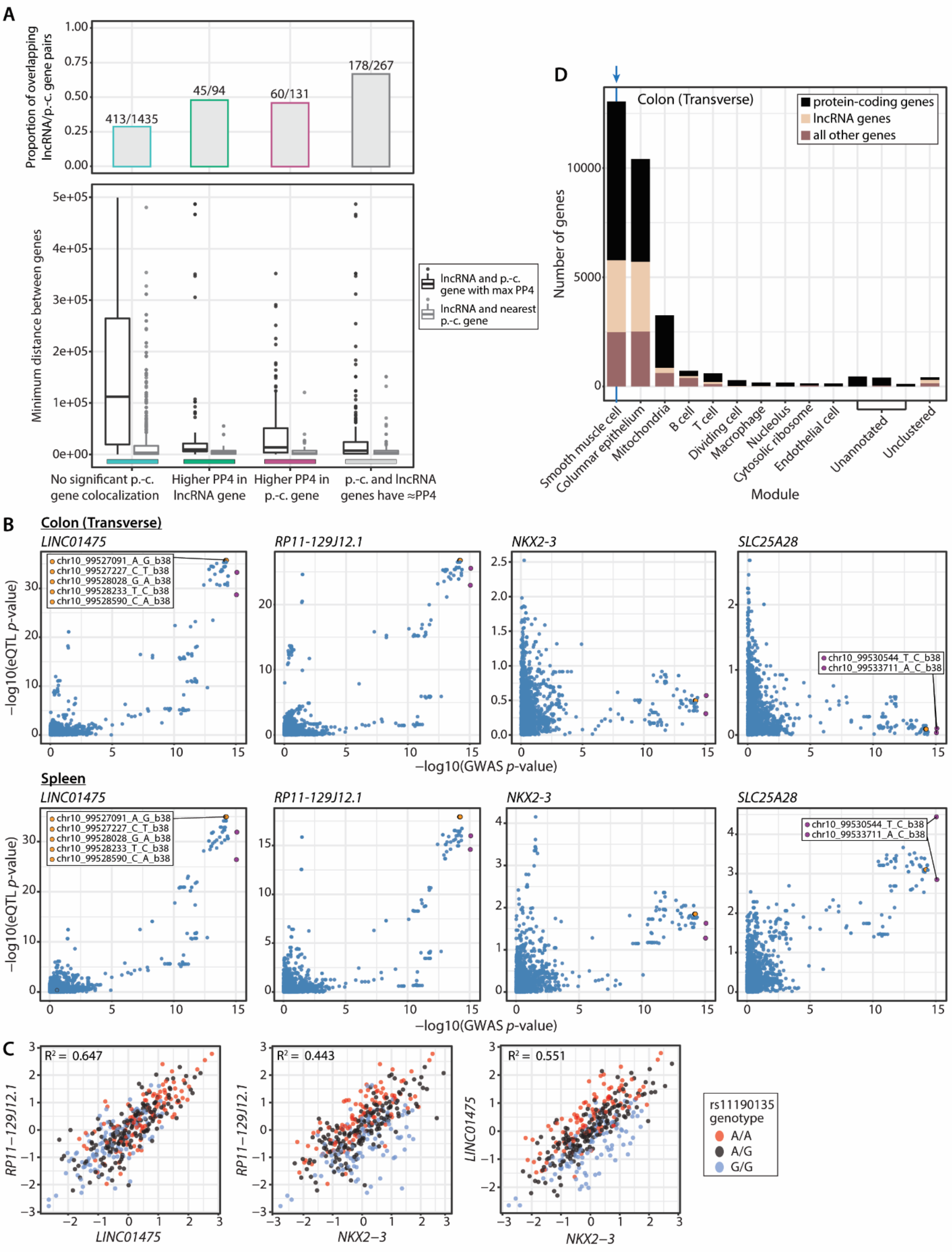
A
Proportion of overlapping lncRNA/p.-c. gene pairs
413/1435
45/94
60/131
178/267
Minimum distance between genes
lncRNA and p.-c. gene with max PP4
lncRNA and nearest p.-c. gene
No significant p.-c. gene colocalization
Higher PP4 in lncRNA gene
Higher PP4 in p.-c. gene
p.-c. and lncRNA genes have ≈PP4
B
Colon (Transverse)
LINC01475
RP11-129J12.1
NKX2-3
SLC25A28
chr10_99527091_A_G_b38
chr10_99527227_C_T_b38
chr10_99528028_G_A_b38
chr10_99528233_T_C_b38
chr10_99528590_C_A_b38
chr10_99530544_T_C_b38
chr10_99533711_A_C_b38
−log10(eQTL p-value)
−log10(GWAS p-value)
Spleen
C
R² = 0.647
R² = 0.443
R² = 0.551
rs11190135 genotype
A/A
A/G
G/G
D
Colon (Transverse)
protein-coding genes
lncRNA genes
all other genes
Number of genes
Smooth muscle cell
Columnar epithelium
Mitochondria
B cell
T cell
Dividing cell
Macrophage
Nucleolus
Cytosolic ribosome
Endothelial cell
Unannotated
Unclustered
Module

**Fig. S9.**

Distance relationships of genes that share significant colocalization events, and further exploration of the significant colocalization of ulcerative colitis and *LINC01475* and *RP11-129J12.1*. (A) Distance between pairs of lncRNA genes with significant colocalization events, and their in-range protein-coding genes with the highest PP4 values. X-axis categories are assigned based on Figure 6A. *(top)* The proportion of lncRNA genes with an overlapping protein-coding gene in which the overlapping protein-coding gene is the one with the highest PP4 value. *(bottom)* The distance between non-overlapping gene pairs. For comparison, the distance between the lncRNA genes and their nearest coding gene is shown as the grey boxplots. (B) Plots of eQTL significance (in the transverse colon and the spleen) versus ulcerative colitis-GWAS significance for *LINC01475*, *RP11-129J12.1*, *NKX2-3*, and *SLC25A28*. The two most significant GWAS variants are colored light blue, and labeled in the *SLC25A28* plots. The most significant eQTL variants for *LINC01475* and *RP11-129J12.1* are colored red, and labeled in the *LINC01475* plot. (C) Normalized expression of *LINC01475*, *RP11-129J12.1*, and *NKX2-3* in transverse colon. Points are colored by the genotype at rs1190135 (chr10_99527091_A_G_b38), one of the most significant eQTLs for *LINC01475* and *RP11-129J12.1*. (D) WGCNA modules of the transverse colon, ordered by annotation and size. The blue arrows indicate the "smooth muscle cell" module to which *LINC01475*, *RP11-129J12.1*, and *NKX2-3* were all assigned.

**Table S1.**

Summary of the 713 lncRNA genes selected for the “lncRNAs with known function” gene group.

**Table S2.**

List of genes with tissue-specific expression for the 28 broad tissues of GTEx. Tissue-specific expression for broad tissues was defined as having TPM $\geq$0.5 in at least 20% of samples of tissues, only for tissues that are grouped in the same broad tissue type.

**Table S3.**

List of tissue-specific eGenes (MashR LFSR <0.05) for the 28 broad tissues of GTEx. To be tissue-specific for broad tissues, an eGene could only be discovered in the tissues that are grouped in the same broad tissue type.

**Table S4.**

Multi-tissue outliers in lincRNA gene expression.

**Table S5.**

Summary of gene pairs that share an eVariant: their expression was either associated with the same eVariant, or with eVariants that were within 500kb of each other with an R2 $\geq$0.85, in the same tissue.

**Table S6.**

Gene-module assignments and connectivity results from weighted gene co-expression network analysis (WGCNA), and module annotations.

**Table S7.**

Summary of the 614 lncRNA/protein-coding gene pairs that shared modules in >12 tissues, as well as sharing at least one multi-tissue gene expression outlier individual.

**Table S8.**

Significant lncRNA gene colocalization events (posterior probability of colocalization PP4 >0.5).

All supplementary tables are available at
https://www.dropbox.com/sh/pkn7rzki5j7yzko/AADbjS3rvzOQUtRgTvnujhoba?dl=0
